# Awake perception is associated with dedicated neuronal assemblies in cerebral cortex

**DOI:** 10.1101/2021.08.31.458322

**Authors:** Anton Filipchuk, Joanna Schwenkgrub, Alain Destexhe, Brice Bathellier

**Affiliations:** Department for Integrative and Computational Neuroscience (ICN),Paris-Saclay Institute of Neuroscience (NeuroPSI), UMR9197 CNRS/University Paris Sud, CNRS, Building 32/33, 1 avenue de la Terrasse, 91190 Gif-sur-Yvette, France; Institut Pasteur, Université de Paris, INSERM, Institut de l’Audition, 63 rue de Charenton, F-75012 Paris, France; Healthy Mind, Station F, 5, parv. Alan Turing, 75013 Paris, France

## Abstract

Neural activity in sensory cortex combines stimulus responses and ongoing activity, but it remains unclear whether they reflect the same underlying dynamics or separate processes. Here we show that during wakefulness, the neuronal assemblies evoked by sounds in the auditory cortex and thalamus are specific to the stimulus and distinct from the assemblies observed in ongoing activity. In contrast, we observed in three different anesthesia, that evoked assemblies are indistinguishable from ongoing assemblies in the cortex. However, they remain distinct in the thalamus. A strong remapping of sensory responses accompanies this dynamical state change produced by anesthesia. Together, these results show that the awake cortex engages dedicated neuronal assemblies in response to sensory inputs, which we suggest is a network correlate of sensory perception.

## Introduction

It has long been noticed that the circuits of sensory areas in the cerebral cortex display intense ongoing activity in the absence of stimuli from their dedicated sensory modality ^1^. The role of this ongoing activity and its relationship to evoked sensory responses remains unclear. Initial observations made under anesthesia in the visual cortex of several mammals ^2–4^ have shown a striking similarity between ongoing activity patterns on the mesoscopic scale and sensory responses, suggesting that ongoing activity could be a form of replay of sensory responses. Similar results have been obtained in the rat ^5,6^ and guinea pig ^7^ primary auditory cortex.

However, recently, recordings in the visual cortex of awake mice have shown that ongoing cortical activity in wakefulness is highly correlated to the level of arousal ^8,9^ and to the animal’s facial motor activity patterns ^10,11^. Arousal-related fluctuations are also seen in the visual thalamus ^12^. Although the direction of causality between behavioral and cortical observables remains to be established, this suggests that ongoing cortical dynamics in the awake state is more than a replay of past sensory activity. In line with this, it was also observed that even if ongoing and evoked activity recruit similar sets of neurons, they correspond to activity patterns that live in orthogonal neuronal dimensions ^10,13^. These conflicting observations across physiological states suggest that anesthesia triggers a profound transformation of neuronal dynamics in the cortical circuits and beyond ^14^. This idea is supported by strong effects of anesthesia on neuronal integration in cortical neurons ^15^. However, the lack of data allowing direct comparison of thalamo-cortical activity patterns at cellular resolution in the same neuronal populations across anesthesia and wakefulness hinders the resolution of this question.

In this study, we imaged on-going and sound-evoked activity in large populations of mouse auditory cortex neurons, as well as axonal terminals from the auditory thalamus, across wakefulness and isoflurane anesthesia. We observed that the cortex generates distinct evoked and on-going cell assemblies during wakefulness which supports an accurate encoding of diverse sounds. In contrast, in anesthesia, on-going and evoked activity patterns became indistinguishable. As a consequence, despite the presence of specific sound responses in the anesthetized state, sound representations were strongly impoverished and were markedly different to the representations observed in the awake state. In thalamic inputs, we observed distinct on-going and evoked assemblies in wakefulness. Under anesthesia however, the two types of assemblies remained different. This indicates a functional disconnection between cortex and its thalamic inputs under anesthesia, whereas the existence of distinct sound-specific and on-going cortical cell assemblies seem to be a signature of awake perception.

## Results

### On-going population events under anesthesia and in wakefulness

Taking advantage of the robustness of GCAMP6s-based^16^ two-photon calcium imaging for the assignment of neuronal activity to identified neurons, we contrasted neural population activity in the auditory cortex of head-fixed mice, in the awake state (**Fig. 1a**) and during light isoflurane anesthesia (**Fig. 1b**). A first 10 min imaging session without auditory stimuli was followed by a 15 min session in which we presented 50 different simple and complex sounds (500ms, **Extended Data Fig. 1**), novel for the animal, each repeated 12 times and delivered in a random order. After sound presentation, another session without auditory stimuli was performed to evaluate the impact of sound presentations on ongoing activity. Then, the same protocol was repeated under anesthesia. 728 +/-180 (474-955) neurons could be imaged simultaneously in cortical layer 2/3 over a 1×1 mm field of view covering about one quarter of the entire auditory cortex. After automated segmentation of the regions-of-interest corresponding to the neurons ^17^, we estimated the time of putative action potentials based on raw fluorescence signal was performed thanks to the MLSpike deconvolution algorithm ^18^ (**Extended Data Fig. 1**). This yielded population activity rasters for well-identified neurons across sessions and physiological states (**Fig. 1c-d, Extended Data Fig. 1**), whose temporal resolution equaled the scan rate of the two-photon microscope (30 Hz).

**Fig 1.**
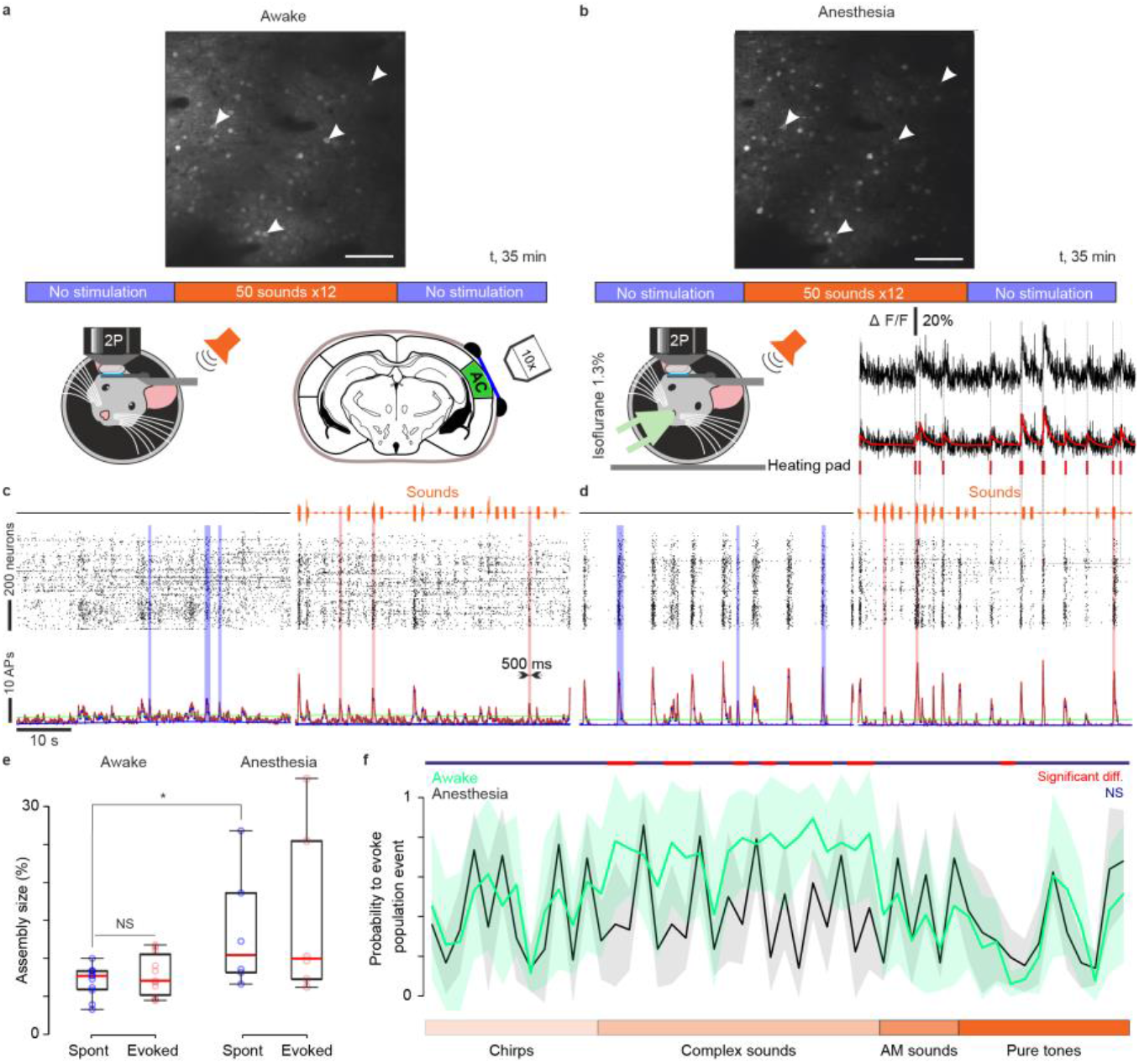
Synchronous population events in the auditory cortex in wakefulness and anesthesia. **(a)** Two-photon Ca^2+^ imaging at 30 Hz sampling rate of up to 1200 layer 2/3 neurons expressing GCamp6s in the awake head-fixed mouse. **Upper panel**: Imaging field-of-view with labeled neurons in awake state showing the standard deviation of fluorescence pixels over a 15 min sound stimulation session flanked by two 10 min long stimulation free periods **(b)** The same recording protocol (**left panel**) and field-of-view (**upper panel**) as in (a) under light isoflurane anesthesia (1.3%). Most neurons are visible in both conditions, demonstrating full stability of the field of view. Weaker or absent labeling in anesthesia reflects neurons that have decreased their activity. Arrows indicate sample neurons. Scale bar is 100 μm. **Right panel:** spikes time estimates (red) were extracted from calcium fluorescence traces (black) using the MLSpike algorithm. **(c)** Population raster plots and population firing rate (30ms bins) during no stimulation (lef t) and stimulation (right) periods. Vertical transparent bars highlight spontaneous (blue) and evoked (pink) population events detected as described in **Fig. S1. (d)**. Same as in (c) for anesthesia. **(e)** The size of ongoing population events increases under anesthesia (p=0.036, n=11 for awake and n=6 for anesthesia, p=0.18 for evoked events, Wilcoxon ranksum test). **(f)** Probability of sounds to evoke population events in the awake state (green traces) and under anesthesia (black trace). Shaded areas indicate standard deviations. The majority of complex sounds significantly decrease their ability to drive population events (p<0.05) when passing from wakefulness to anesthesia (Wilcoxon Rank Sum Test for **e**-**f**).

In line with previous observations ^5,19–21^, inspection of the raster plots and of the instantaneous population firing curves revealed short synchronous population activity events, which occurred both in the anesthetized and awake states but seemed more contrasted and stereotypical under anesthesia (**Fig. 1c-d, Extended Data Fig. 1**). We extracted these events by applying a baseline corrected ^22^ threshold to the population firing rate above which a synchronous event could not be explained by the fluctuations of summed independent Poisson processes (**Fig. 1c-d, Extended Data Fig. 1**). Population events were short but of variable durations (327+/-131ms, n=11 mice in awake state, 414+/-237 ms, n=6 mice under anesthesia) and appeared both during the stimulation-free (0.42+/-0.1 Hz, n=11 mice in awake state, 0.51+/-0.11 Hz, n=6 mice under anesthesia) and sound delivery protocols (**Fig. 1c**). The percentage of neurons recruited in each population event increased under anesthesia (**Fig. 1e**). Relying on the same detection criterion, the probability that a population event appeared during sound presentation was 52+/-20%, (n=11 mice) in the awake state and 41+/-8% (n=6 mice) under anesthesia, with a large disparity across sounds. Some sounds drove detectable population events on almost every trial, others did not (**Fig. 1f**). The probability to evoke a population event significantly changed for many sounds between wakefulness and anesthesia (**Fig. 1f**) in line with previous reports ^23,24^, suggesting that anesthesia reorganizes cortical responses.

### Indistinguishable on-going and evoked cortical activity patterns under anesthesia

To investigate more precisely this reorganization, we then asked if events observed in the absence of stimuli resemble the activity generated by sensory stimuli, and in particular if similar assemblies of neurons are recruited in both cases. For on-going activity, we identified each detected population event to the assembly of neurons that fired at least one putative action potential during the event. To avoid unnecessary thresholding effects, for sound-evoked activity, the assembly of responsive neurons was the neurons that fired at least one putative action potential during sound presentation (response time window: 0 to 500ms after sound onset). Similarity between assemblies was measured based on the correlation between binary population vectors (entry 1 for neurons belonging to the assembly and 0 otherwise), which had the length of the entire neuronal population. Then we used a simple approach to account for the intrinsic variability of neural responses and noise introduced by spike estimation errors ^18^. We performed hierarchical clustering to organize all ongoing assemblies in groups of similar patterns (independently for ongoing activity before and after stimulation) and displayed the matrix of pairwise similarity between individual ongoing assemblies and single trial responses to all tested sounds (**Extended Data Fig. 1, Fig. 2 a,b**). Visual inspection of sample matrices for pairs of sounds and ongoing assembly clusters indicated overall a low similarity between ongoing assemblies and evoked sound responses in the awake state contrasting with high similarity levels in anesthesia in the same neuronal population (**Fig. 2a,b**). To quantify this, we measured, for each sound and imaging session, the mean similarity between individual evoked responses and the assemblies of the most similar ongoing assembly cluster. To evaluate to which extent, similarity is limited by the variability of responses or spontaneous pattern, we defined the reproducibility of evoked responses as the mean population activity correlation across repetitions of the same stimulus. Reproducibility of spontaneous clusters was defined as the mean correlation between the assemblies within a cluster. Plotting similarity against the mean of spontaneous and evoked assembly reproducibility, it became evident that similarity was below reproducibility levels in the awake state whether ongoing assemblies were taken before (**Fig. 2c**) or after (**Fig. 2d**) sound presentations. This result was robust to changes in clustering parameters (**Extended Data Fig. 2**), and cortical depth (**Extended Data Fig. 3**). It also held for assemblies observed in between sound stimulations (**Extended Data Fig. 4**), and when on-going assemblies were compared to sound responses clustered in the same way (**Extended Data Fig. 2**). Hence, evoked responses and ongoing assemblies correspond to distinct population activity patterns in awake freely listening mice.

**Fig 2.**
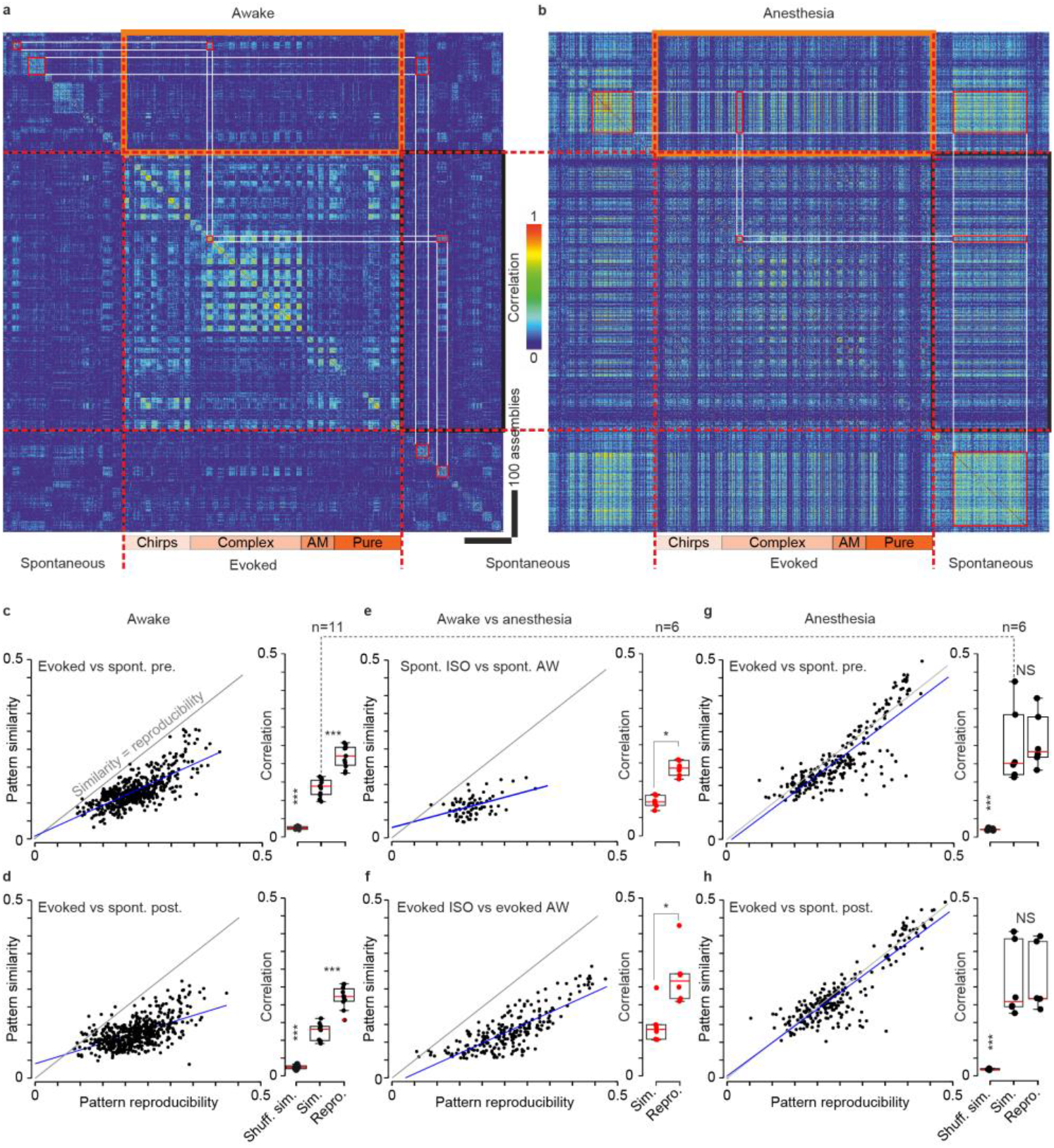
Ongoing assemblies and sound-evoked responses differ in the awake state but overlap under anesthesia. **(a)** For an example recording session, Pearson’s correlation matrix between spontaneous assemblies sorted by hierarchical clustering and single trial sound response patterns (whether or not a population event was detected), sorted sound by sound (12 trials/sound, sound order indicated below). Clustering is done independently in pre-and post-stimulation periods. Lower correlation inside black and orange frames (similarity) compared to correlations along the diagonal (reproducibility) indicate that spontaneous and evoked patterns are different. **(b)** Correlation matrix under anesthesia for the same neuronal population as in **a**. Similar correlation in black and orange frames (similarity) and in the squares along the diagonal (reproducibility) indicate that spontaneous and evoked assemblies are highly similar. **(c-h)** Relation between reproducibility (abscissa) and similarity (ordinate) of sound-evoked and spontaneous patterns for all sounds and sessions. Statistics across sessions are given on the right-hand-side histograms (**c**: p=0.001, p=0.001, **d**: p=0.001, p=0.001, Paired Wilcoxon Signed Rank Test, n=11 mice). Spontaneous and evoked patterns can be considered dissimilar if their reproducibility is significantly larger than their similarity (grey: line for the equality, dark blue: data trend). A significant difference between spontaneous and evoked patterns is seen in awake mice (**c-d**), but not under anesthesia (**g-h**, p=0.44, p=0.56, Paired Wilcoxon Signed Rank Test, n=6 mice) where the similarity increased significantly (dashed line, **c vs g**, p=0.0003, Wilcoxon Ranksum test). Evoked and spontaneous population activity patterns under anesthesia are different from corresponding ones in the awake state (**e-f**, p=0.03, p=0.03 Paired Wilcoxon Signed Rank Test, n=6 mice)

However, in anesthesia a profound reorganization of cortical dynamics was observed. First, ongoing assemblies and sound responses seen under anesthesia were clearly distinct from those seen in the awake state (**Figs. 2e-f, Extended Data Fig. 5**). Second, we observed that evoked responses and ongoing assemblies are highly similar under isoflurane anesthesia, as measured through the equal similarity and reproducibility levels of particular sound responses and with at least one cluster of spontaneous assemblies (**Figs. 2g-h, Extended Data Fig. 5**).

Overall, the massive transformation of assemblies from wakefulness to anesthesia can be also be qualitatively summarized by plotting the localisation of on-going assemblies in the neuronal state space after dimensionality reduction (i.e. in the space of the first three principal components of dataset including all assemblies and response, **Fig. 3**). In this format, it becomes evident that assemblies observed in the awake state are essentially distinct from assemblies observed under anesthesia. Moreover, under anesthesia, sound responses and ongoing assemblies span the same region of the neuronal state space, while in wakefulness they clearly span different regions (**Fig. 3**). In the awake state, spontaneous and evoked activity can thus be easily distinguished, while under anesthesia, a sound response leads to population activity patterns that are extremely similar to ongoing activity. Therefore, sound responses under isoflurane are much harder to interpret as a stimulus to be perceived in downstream targets of the auditory cortex, which could potentially contribute to the lack of perception despite the existence of cortical responses.

**Figure 3.**
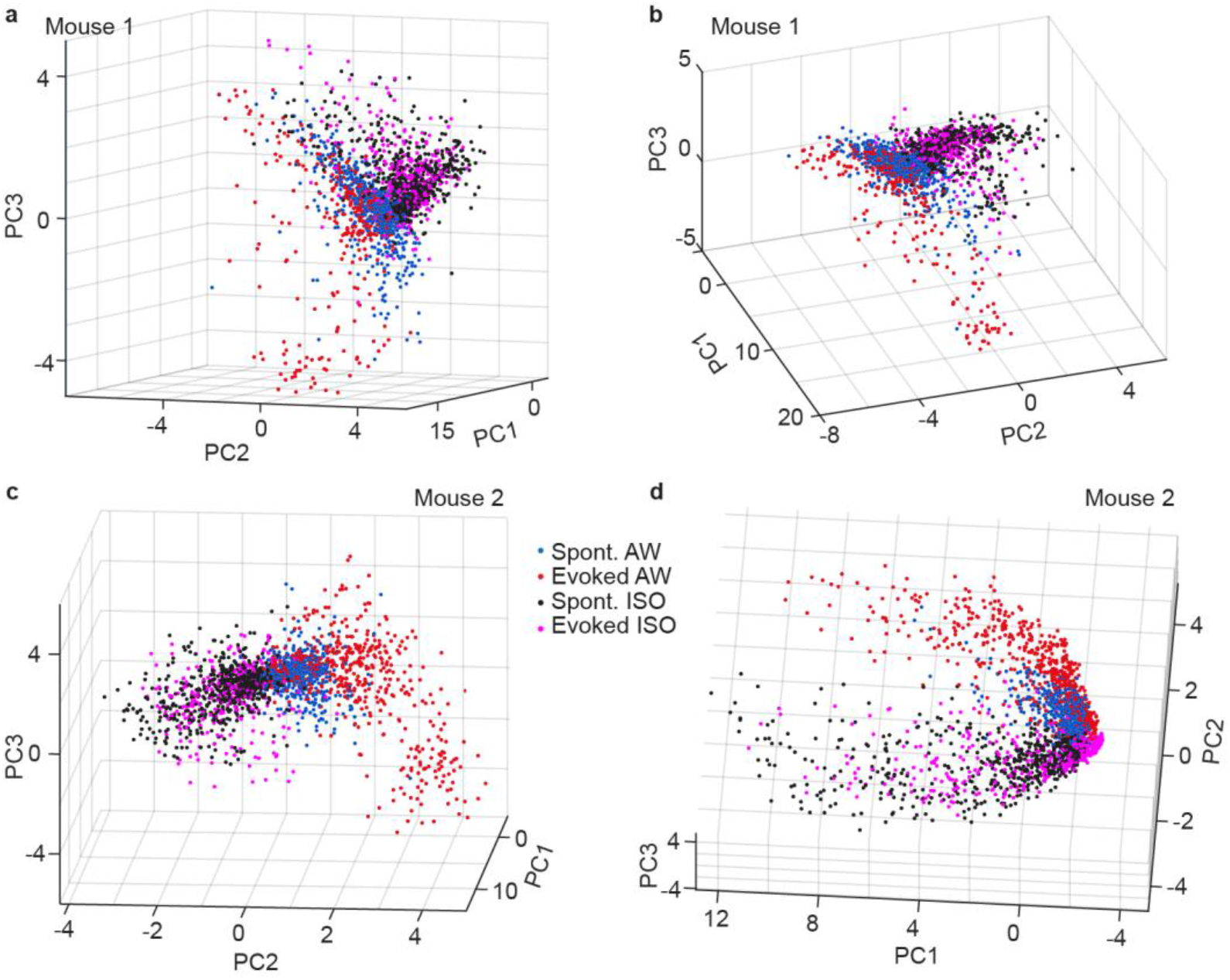
Assemblies of the awake and anesthetized states span different regions of the population state space. **(a)** Plot of the localisation of ongoing and evoked assemblies in the state in the three-dimensional space defined by the first three principal components of the dataset including all on-going and evoked assemblies, for one representative recording sample. Anesthesia: isoflurane. **(b)** Same as (a) but with a different three-dimensional view angle. **(c-d)** Same as (a-b) for another sample recording in a different animal. Anesthesia: isoflurane. Color code: magenta = evoked isoflurane, red = evoked awake, black = spontaneous isoflurane, blue = spontaneous awake.

In order to further probe this idea, we repeated the experiment with other anesthetics that have different modes of action. Following the same neuronal populations in the auditory cortex, before and after anesthesia induction with subcutaneous injection of the classical mix of 50 mg/kg ketamine and 1 mg/kg medetomidine (KM), we observed that evoked responses and ongoing assemblies become extremely similar with anesthesia like with isoflurane (**Extended Data Fig. 6**). This is remarkable as KM, unlike isoflurane, is a dissociative anesthetic that targets MNDA receptors. In the mouse, it also leads to faster cortical rhythms than isoflurane as we could measure for the neuropil signal, a proxy of ECoG signals ^25^ (**Extended Data Fig. 7**). Often used in auditory neurophysiology experiments, KM provides long-lasting anesthesia in mice and can produce deep anesthesia. We therefore also repeated our experiments with the lighter dissociative anesthetic Zoletil®, a 50-50% mix of tiletamine and zolazepam (70 mg/kg), that maintains cardiorespiratory function at a high level. This led to the same results (**Extended Data Fig. 6**) despite shorter anesthesia duration (∼typically 40 min) and a different associated brain rhythm (**Extended Data Fig. 7**). Note that in all cases, we could verify in all three anesthetics that population activity followed global up and down states, which is a clear sign of anesthesia (**Extended Data Fig. 7**).

### Impoverished cortical sound representations under anesthesia

These results also suggest that assemblies in on-going activity and in sensory responses emerge from a more stereotypical process in anesthesia, while in the awake state, the auditory cortex develops a richer repertoire of activity patterns which carries more information, in particular about sounds. In order to investigate this, we reordered all neurons imaged in both conditions based on a hierarchical clustering of their trial-averaged response signatures to our 50 sounds in the awake state (**Fig. 4a**). The number of clusters was chosen to be close to the maximum dimensionality of the pool of response signatures which is limited by the number of sounds ^26^. This revealed a variety of response signatures that corresponded to groups of neurons of different sizes as plotted in the heatmap of **Fig. 4a**. When plotting with the same order of the neurons the heatmap corresponding to sound response signatures during anesthesia, it appeared that many of the neurons displaying responses that were specific to fewer sounds in the awake state responded under anesthesia either with an absence of responses or with a less specific and more stereotypical response signature (**Fig. 4b**). This was also clearly visible when plotting the mean response signatures for awake and anesthetized states for a few representative clusters (**Fig. 4c**). Several clusters showed highly significant changes of their responses. For a few of them (e.g. cluster 1 and 16), this corresponded to minor changes in response magnitudes. However, for many clusters, anesthesia drastically changed their mean response signature, often due to the disappearance of some sparse, specific responses (**Fig. 4d**). This suggested that the sound representation was strongly impoverished by anesthesia. Interestingly, the few clusters with response signatures that were preserved under anesthesia (representing 43% of neurons), suggest the existence of more robust response modes (e.g. **Fig. 4a-c** clusters 1, 2 and 16). To quantify the impoverishment of sound-evoked assemblies patterns we evaluated the information carried by sound-evoked population responses in each imaging session using a cross-validated template matching classification algorithm. Corroborating our qualitative observations, overall sound decoding performance drastically dropped under anesthesia (41 +/- 10 % in awake; 7 +/-3%), without reaching the 2% chance level (**Fig. 4e**). Moreover, the structure of prediction errors was strongly modified between the awake and anesthesia state (**Extended Data Fig. 5)** corroborating the profound change of sound representations at population scale under anesthesia. One reason for the lower sound information despite continued sound responses, is that single neurons tended to be less specific of particular sounds under isoflurane anesthesia (**Fig. 4d**). This could be quantified with a generic lifetime sparseness measure, the kurtosis of the response distribution ^27^, across a wide range of clustering sizes (**Fig. 4f**). Interestingly, sparser clusters were more affected by anesthesia (**Fig. 4g**), indicating that, even if anesthesia spares a large fraction of sound responses, it tends to abolish the most specific ones. Therefore, anesthesia produced a strong impoverishment of sound representations which preserved only a small fraction of the sound information that was present in the same cortical neuron population in the awake state.

**Figure 4.**
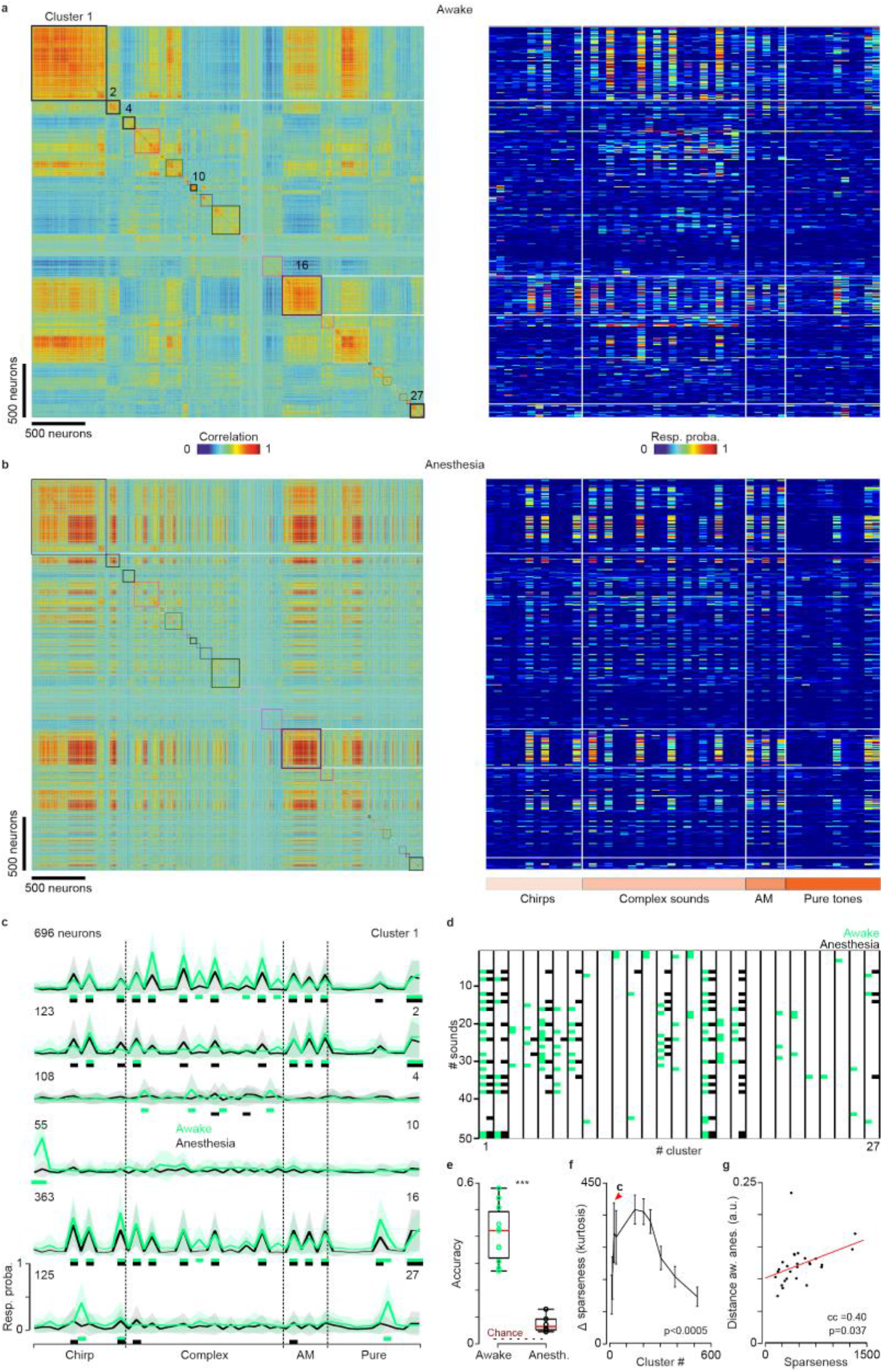
Modification of sound tuning between wakefulness and anesthesia. **(a) Left panel** : Matrix of response profile correlation for the 3641 (6 mice) recorded neurons in the awake state. The neurons are clustered according to the similarity of their responses (metric: Pearson correlation coefficient between response profiles based on probabilities to respond to any out of 50 sounds). **Right panel:** Response probability for all recorded neurons organized with the same clustering as for the matrix in the left panel (colors in the upper band indicate the type of sound). **(b)** Same neurons as in (a) under anesthesia (neurons and response profiles are ordered according to the clustering made in the awake state). **(c)** Mean response profiles of auditory cortex neurons in awake state (green) and under anesthesia (black) plotted as the mean firing rate of the 6 sample clusters labeled in (a). Green and black rectangles indicate responses significantly different from 0. **(d)** For the 27 neuronal clusters identified in **a**., brown squares indicate the sound which produced a significant response in the neurons of the cluster in awake (green) or in anesthesia (black) assessed with a multicomparison 1-way Anova between the actual responses and a shuffled surrogate of the responses. **(e)** Average prediction accuracy for all sounds mixed in awake vs anesthetized state (n=11 and 6, p=0.0002, Wilcoxon Rank Sum Test). **(f)** Difference of sparseness of cluster responses to sounds (i.e. kurtosis of response distribution) between awake and anesthetized states, as a function of the number of the granularity of the clustering algorithm (i.e. final number of clusters). Positive values indicate higher sparseness in the awake state. All measures are significantly larger than zero (Wilcoxon Signed Rank test, p<0.0005, error bars = SEM). (**g**) Difference between sound responses in awake and in anesthesia as a function of response sparseness for the 27 clusters selected in **c**.

### Distinct spontaneous and evoked thalamic activity patterns in wakefulness and under anesthesia

To evaluate the contribution of thalamic inputs in the reorganization of cortical activity, we imaged thalamic boutons in the auditory cortex labeled with GCamP6s through stereotaxic AAV virus injections in the primary and secondary auditory thalamus (**Fig. 5a**) ^28,29^. Signals from individual thalamic axons and boutons were extracted using the same automated methods as for somatic fluorescence, applied to a smaller field of view in layer 1 of the auditory cortex. Using the MLSpike algorithm we obtained estimates of the spike trains (**Fig. 5a**) underlying thalamo-cortical synaptic release in cortex. As in the cortex, population activity in the thalamus displayed short synchronous population events which more clearly departed from baseline population activity under anesthesia (**Fig. 5b-c**). Measuring population vector correlation as previously (**Fig. 5d,e**), we observed that ongoing assemblies were significantly different from evoked activity in the awake state in the thalamic output as in the cortex (**Fig. 5f,g**). Thus, the thalamic input provides separate information streams based on which the cortex can build distinct assemblies in its ongoing and evoked activity.

**Figure 5.**
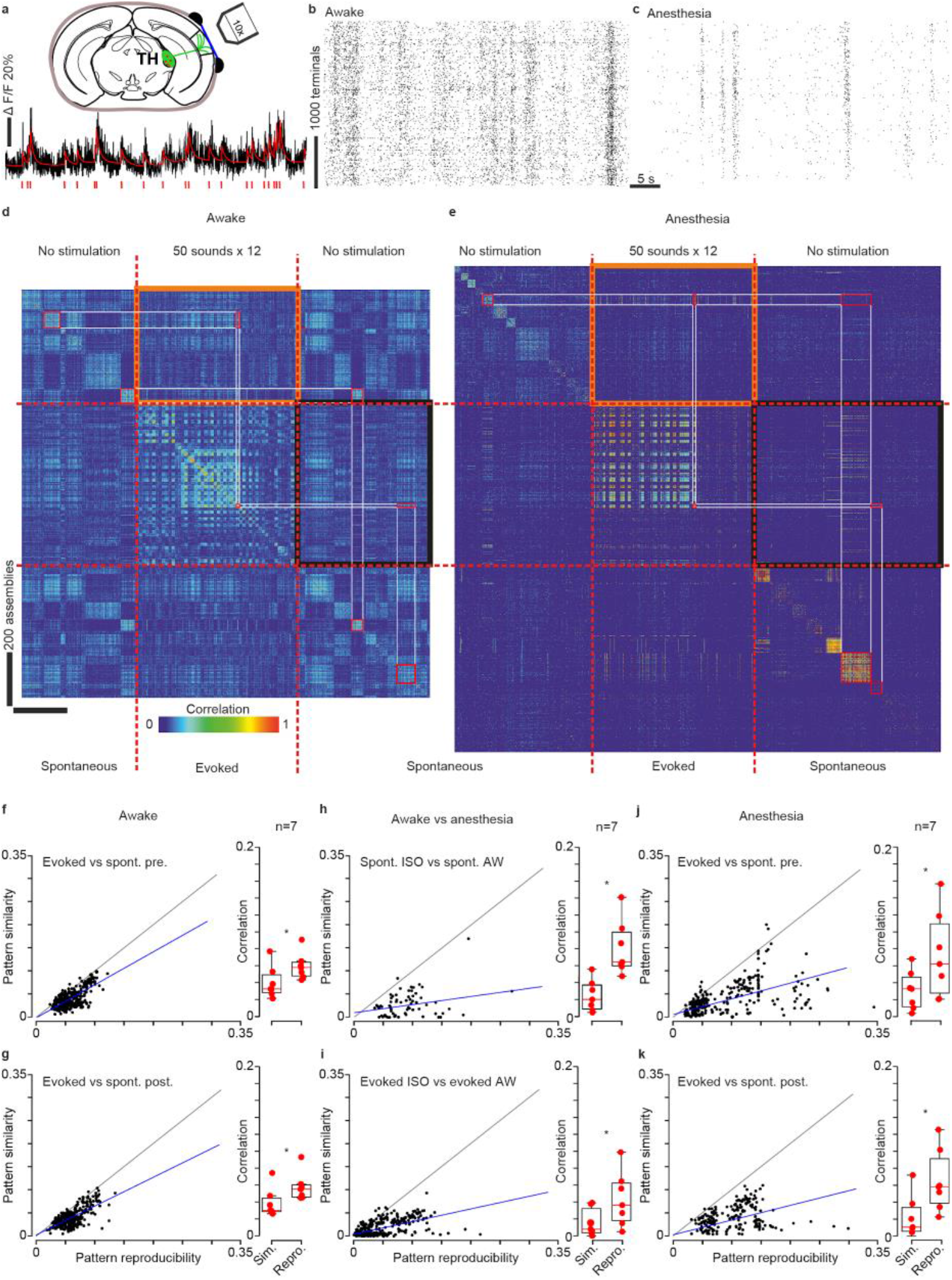
Ongoing and evoked population activity patterns in auditory thalamus differ both in the awake and anesthetized states. **(a)** Two-photon Ca^2+^ imaging at 30 Hz sampling rate of up to 4000 thalamo-cortical terminals expressing GCamp6s (AAV1 viral injection into the thalamus) in the AC layer 1 of awake head-fixed mouse. Below: example of thalamo-cortical terminal calcium trace (black) with spikes time estimates (red) using the MLSpike algorithm. **(b)** Population raster plot in awake state where each line represents spontaneous spiking pattern of a thalamo-cortical terminal. **(c)** Same under light isoflurane anesthesia (1.3%). **(d)** For an example recording session, Pearson’s correlation matrix between spontaneous assemblies of thalamo-cortical terminals sorted by hierarchical clustering and single trial sound response patterns (whether or not a population event was detected), sorted sound by sound (12 trials/sound) **(e)** Same as **d** under anesthesia. Lower correlation inside black and orange frames (similarity) compared to correlations along the diagonal (reproducibility) indicate that spontaneous and evoked patterns are different. **(f-k)** Relation between reproducibility (abscissa) and similarity (ordinate) of sound-evoked and spontaneous patterns for all sounds and sessions. Statistics across sessions are given on the right-hand-side histograms (**f-k**: p=0.016, Paired Wilcoxon Signed Rank Test, n=7 mice). Spontaneous and evoked patterns are dissimilar in both awake (**f-g**) and anesthetized states (**j-k**) (gray: line of equality, dark blue: data trend) and neither of them keep similitude passing from one state to another (**h-j)**.

Interestingly, under anesthesia, neural assemblies in the thalamic input were also profoundly reshaped as compared to assemblies seen in the awake state (**Fig. 5h, i**). This involved changes in the tuning of individual neurons to sounds, parallel to those observed in the cortex (**Extended Data Fig. 8**). During anesthesia but not during wakefulness, we often observed thalamic activity patterns were usually different across our two spontaneous activity recording sessions (before vs after sound stimulation, **Fig. 5e**). This may reflect the fluctuations in the depth of anesthesia typically observed in such narcosis conditions. However, unlike what we observed in cortical neurons, ongoing assemblies observed before or after sound presentation in thalamic fibers activity were clearly different from evoked activity patterns, as seen from reproducibility levels that were at least twice larger than similarity levels (**Fig. 5j, k**). Therefore, during anesthesia, the thalamus sends sensory driven and ongoing activity inputs that are different, but cortical activity does not take this difference into account.

### Specific cell populations for evoked and spontaneous activity patterns

To find further evidence for this observation, we analyzed single neuron properties in cortical activity, seeking to identify distinctive functional markers of ongoing and evoked activity. We measured for each neuron the probability to emit at least one putative action potential during a spontaneous assembly and during presentation of a sound (500 ms after onset). In the awake state, the scattering of probabilities was broad (**Fig. 6a**). Substantial fractions of neurons were more specific to ongoing assemblies (∼1/6) or more specific to sound responses (∼1/6, **Fig. 6a**). Remaining neurons displayed similar probabilities to participate in ongoing assemblies or evoked responses (∼2/3, **Fig. 6a**). Specific neurons were large contributors to the identity and robustness of on-going assemblies and sound responses. Indeed, if specific neurons were removed by retaining neurons for which the absolute difference between ongoing and evoked probabilities was below one mean absolute deviation of the full distribution, population vector reproducibility levels drastically dropped both for ongoing assemblies and sound responses (**Fig. 6b**), although some information about sounds remained (**Extended Data Fig. 9**). Corroborating this, when non-specific neurons were discarded, population vector reproducibility levels were boosted (**Fig. 6c-e**). Also, if only spontaneous assembly neurons were retained, reproducible and sound-specific sound response patterns disappeared (**Fig. 6d**), leading to a severe drop in sound decoding (**Extended Data Fig. 9**). This indicates that even if no neuron is fully specific of on-going or evoked activity as previously observed in the visual cortex ^10^, small and distinct neuronal subpopulations are specialized in carrying sound specific information *versus* forming different on-going activity motifs.

**Figure 6.**
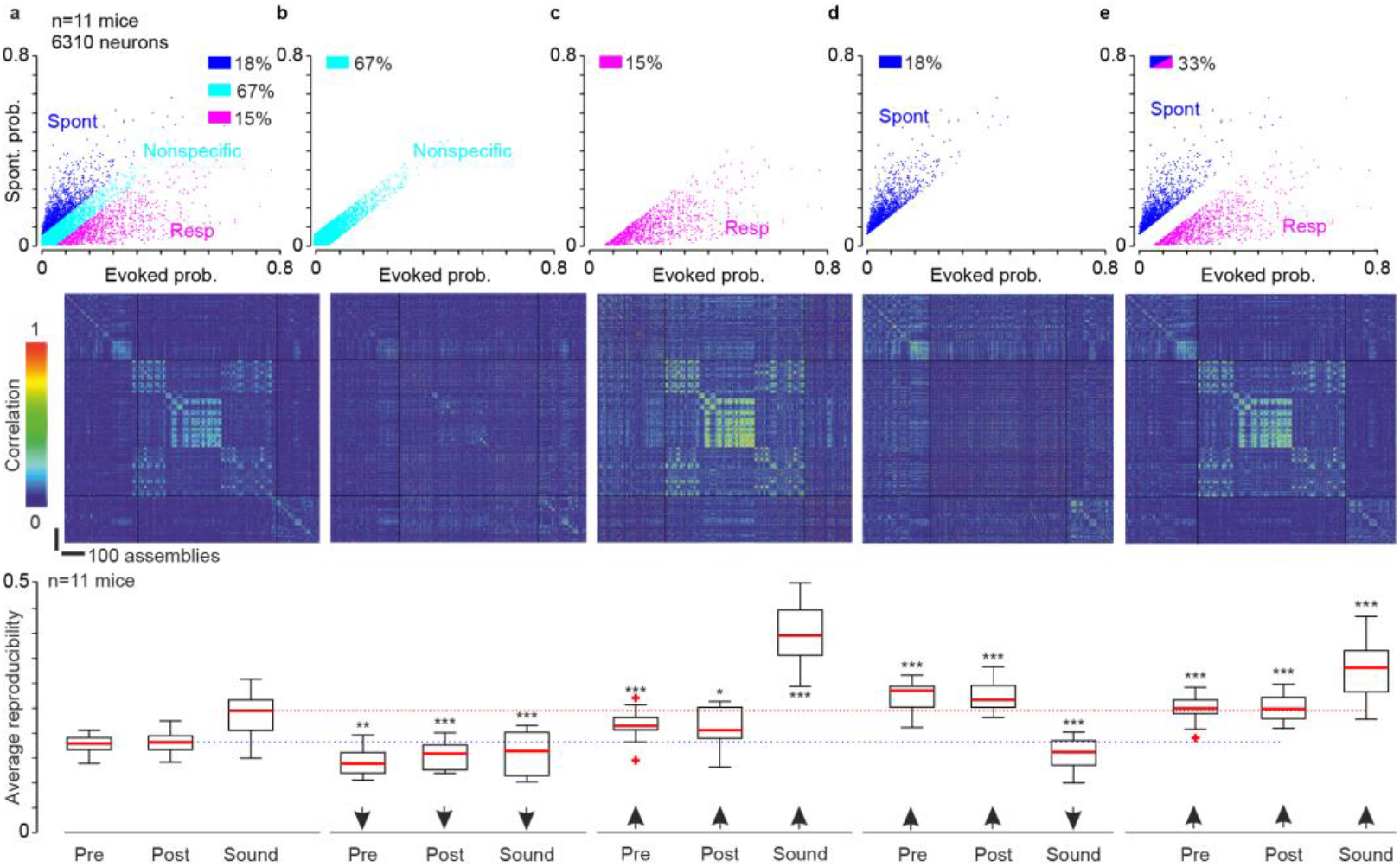
Specificity of evoked patterns rely on a subset of neurons in the awake state. **(a) Top:** Plot of the probability to respond to any sound versus the probability to be recruited in an on-going event for 6310 neurons over 11 mice. The color code indicates whether the neuron prefers on-going (blue) or evoked (magenta) events or nonspecifically participates in both types of events (cyan, boundaries: +/- 1 mean absolute deviation of the probability difference). **Middle**: For a sample session, Pearson’s correlation matrix computed with all available neurons. **Bottom**: Average sound and spontaneous event cluster reproducibility (**b-e**: p=0.002/0.001/0.001 p=0.001/0.02/0.001, p=0.001/0.001/0.001, p=0.001/0.001/0.001 Paired Wilcoxon Signed Rank Test, n=11 mice). **(b)**. Same as **a**, but correlation matrix and reproducibility (mean correlation across assemblies of the same spontaneous cluster or of the same sound) are calculated with nonspecific neurons only (67%). **(c)**. Same as **a**, but for sound responsive neurons only (15%). **(d)**. Same as **a**, but only for neurons preferring on-going events (18%). **(e)**. Same as **a** but for all except nonspecific neurons (33%).

Tracking non-specific, sound specific and ongoing assembly specific subpopulations during anesthesia further uncovered the profound transformation of cortical dynamics. Plotting the probability to fire in an ongoing assembly against the probability to fire in a sound-evoked response indicated that the two specific neuronal subpopulations were strongly redistributed under anesthesia (**Fig. 7a, Extended Data Fig. 9**). To validate this observation beyond our measure of participation to cell assemblies, we plotted the mean firing rate of each of the three subpopulations over the sound stimulation and stimulation-free phases of our protocol prior to assembly identification. In line with its participation in sound evoked assemblies in the awake state, the sound specific population displayed increased activity during sound stimulation compared to the stimulus-free phases. Likewise, the ongoing assembly specific population is more active in the stimulation-free period. When plotting the activity of the same subpopulations under anesthesia, the modulations disappeared (**Fig. 7b, Extended Data Fig. 9**), validating the idea that single cell properties are massively reassigned under anesthesia.

**Figure 7.**
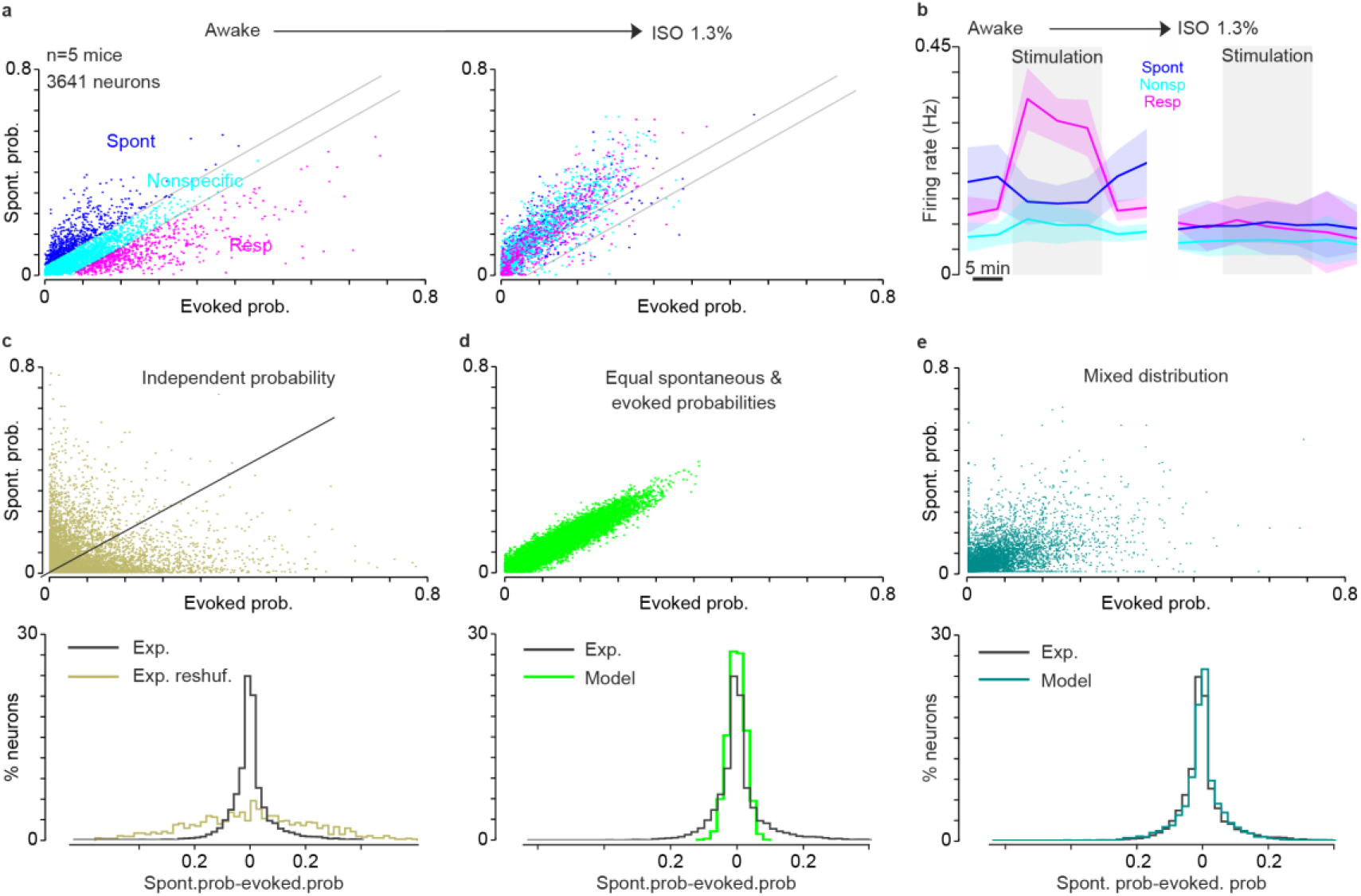
Specificity of cortical neurons for evoked or ongoing activity vanishes in anesthesia. **(a)**. Plot of the probability to be recruited in an on-going event (spont. prob.) plotted against the probability to respond to any sound (evoked prob.) for 3641 neurons over 5 mice in awake (left) and under anesthesia (right). The color code is defined as in A for the awake state. Under anesthesia, the three color-coded populations converge to a single group with strongly correlated probability of activation in spontaneous and evoked events. **(b)**. Time course of the mean firing rate profiles for each of the 3 groups of neurons in awake (left) and under anesthesia (right). Under anesthesia, specificity to sound and ongoing activity disappears. **(c) Top:** same as **a**, but probabilities are reshuffled along each axis to show the expected probability distribution for independent participations to on-going and evoked events. **Bottom**: corresponding distribution of probabilities difference (on-going -evoked, 6310 neurons, over 11 mice). Experimental (black), reshuffled (gray green). **(d)** Probability distributions for 6310 surrogate neurons with equal participation probabilities to on-going and evoked events. **(e)** Probability distributions for 6310 surrogate neurons whose probabilities to participate in spontaneous and evoked activity are correlated but not equal (i.e. the difference between spontaneous and evoked probabilities is drawn from a continuous Gaussian distribution).

To more precisely interpret the distribution of single cell activity in wakefulness, we simulated three different hypotheses about the relationship between the probability to be recruited in a sound-evoked response and the probability to be recruited in an ongoing assembly. The first hypothesis we considered was that evoked and ongoing probabilities are independent. To estimate the probability distribution that would result from this hypothesis we reshuffled all probabilities across cells. This generated a much broader distribution of ongoing and evoked probabilities across cells than the one actually observed (**Fig. 7c**). This indicates that there exists significant correlation between engagement probabilities in sound-evoked responses and on-going assemblies both in the awake state and under anesthesia. The second hypothesis was that ongoing and evoked probabilities are equal and that the observed distribution arises from probability estimation errors. This hypothesis led to a much narrower distribution of probabilities than observed in wakefulness but also under anesthesia (**Fig. 7d** vs **7a**). The mean absolute distance of data points from the linear regression line fitted to spontaneous and evoked probabilities was 0.010 if we suppose equal recruitment probabilities (Fig. 7d), whereas it was 0.032 and 0.018 in the data for the awake and anesthetized states respectively. The differences between all three measures were highly significant (p<0.001 bootstrap test with 1000 resamplings) indicating that despite clear correlations between the probabilities to engage in spontaneous and evoked events, neurons also tend to specialized for one or the other, and more so in the awake than in the anesthetized state.

In line with this, the data were much better approximated if we modeled the probabilities to engage in a spontaneous or an evoked assembly as the sum of a common and independent probability, chosen randomly within continuous distributions (**Fig. 7e**). Together, this analysis shows that, under anesthesia, the recruitment of a cortical neuron in a sound response is more strongly determined by its probability to be recruited in a spontaneous event, leaving less freedom to encode sound information. In the awake state, the two probabilities are more dissociated, opening much larger possibilities to encode sound information.

## Discussion

Our data show that sensory inputs and ongoing activity engage distinct neuronal assemblies in cortex during wakefulness, that during anesthesia sensory responses generate assemblies that also appear in on-going activity. This observation reconciles contradicting findings obtained in the two different states. In anesthetized animals a recurrence of evoked responses in ongoing neuronal assemblies had been observed at mesoscopic and cellular resolutions ^2,5^. This contrasted with recent reports in awake animals indicating that on-going activity is mostly orthogonal to evoked responses ^10,11^. We provide here a comparison of the spontaneous and evoked cell assemblies in both states, together with thalamic activity. We found that although thalamic inputs are distinct, the anesthetized cortex engages stereotyped cell assemblies, already present in spontaneous activity. In contrast, sound-specific cell assemblies are evoked in awake conditions, when sounds are perceived by the animal.

One main observation (**Fig. 2**) is that the cell assemblies evoked in the anesthetized cortex appear to engage cell assemblies already present in the spontaneous activity, where both spontaneous and evoked activity stem from the same restricted set of cortical cell assemblies, while much richer sets of assemblies are seen in the awake cortex. This restriction of the dynamics during anesthesia may explain previous findings indicating a low dimensionality of sound responses in the anesthetized auditory cortex ^19,30^. Interestingly, if sensory information collapses under anesthesia (**Fig. 4**), it does not fully disappear, as indicated by sound classification performance clearly above chance levels and by the fact that almost half of the neurons retain their response profile. This is in line with the common observation that stimulus specific patterns still exist under anesthesia ^23^. Although we could not directly assess this aspect, previous reports indicate that activity patterns in anesthesia follow well-known functional maps like, for example contour orientation maps in the visual cortex of carnivores ^2,31^ or the tonotopic map in the auditory cortex ^7,32^. These maps correspond to anatomically hardwired circuits ^33,34^ which may constrain the spatial extent of the stereotypical population events that dominate cortical dynamics during anesthesia (**Fig. 1**). In humans, resting state activity also reduces its complexity and follows large scale anatomical connectivity structures in anesthesia ^35^. Our results further indicate that while preserving these large scale spatial features and the activity of a large number of individual neurons, anesthesia abolishes responses carrying more precise and more sparsely encoded information about the stimulus (**Fig. 4**). Comparisons of anesthetized and awake datasets from different experiments have suggested the absence of important features of auditory responses in the cortex such as offset ^36^ or sustained responses ^37,38^ in anesthesia. Here we could directly show that, in anesthesia, many neurons change or lose the specific sound response properties that they had during wakefulness. Although it has been reported that pure tone response properties are affected by anesthesia ^23,39^, the use of diverse complex sounds in our study may have magnified the effects of anesthesia, by highlighting more diluted aspects of the auditory code ^40^ which seem also more sensitive to anesthesia (**Fig. 4**).

Thanks to large scale imaging at single cell resolution, our results revealed a profound change in the cortical and thalamic population dynamics at a mesoscopic scale with anesthesia, while activity at the individual neuron level is partially preserved (**Figs. 1-4**). This indicates that anesthesia induces network-scale effects across cortical layers (**Extended Data Fig. 3**) and thalamo-cortical connections (**Fig. 5**). How anesthetics such as isoflurane which potentiates inhibition^41^ lead to this profound change of dynamics remains a challenge for neural network modeling. The fact that unlike the cortex, the thalamus generates different assemblies for on-going and evoked activity in anesthesia indicates some degree of functional disconnection between the two structures, as the anesthetized cortex seems to ignore a difference present in its inputs. This indicates that the emergence of distinct cortical assemblies in awake conditions is of cortical origin.

The collapse of information through a dynamical change has been readily proposed as a mechanism of the loss of consciousness under anesthesia along with a loss of functional connectivity ^14,42,43^. Our results provide a strong quantitative support for this theory and suggest that the ability for the sensory cortex to form stimulus-specific neuronal assemblies that encode precise information about the sensory input could be a marker of wakefulness with respect to anesthesia. Because sensory input just evokes patterns of spontaneous activity, one may assume that propagation of sensory information to other cortical areas will be limited, in agreement with findings in human anesthesia ^44,45^. This may also explain the loss of perception during anesthesia, and our results show that this loss already occurs in the primary sensory cortex. Finally, our results appear compatible with the idea that in the awake cortex, the new assemblies formed by sensory inputs will propagate across the brain, but the mechanisms underlying this selective propagation are presently unknown, although previous models have proposed that asynchronous states may be the substrate of this selective propagation ^46^.

## Methods

### Animals and surgery

We used C57bl6 male and female 8-16 weeks mice. Animals were housed 1–4 animals per cage, in normal light/dark cycle (12 h/12 h). All procedures were in accordance with protocols approved by the French Ethical Committee (authorization 00275.01).

To allow chronic unilateral access to auditory cortex (AC) for 2-photon calcium imaging, a cranial window was incorporated into the skull and a metal post for head fixation was implanted on the contralateral side of the craniotomy. Surgery was performed in 4 to 6 week old mice placed on a thermal blanket under anesthesia using a mix of ketamine (80 mg/kg, Ketasol) and medetomidine (1 mg/kg, Domitor, antagonized with atipamezole -Antisedan, Orion pharma – at the end of the surgery). The eyes were covered using Ocry gel (TVM Lab) and Xylocaine 20mg/ml (Aspen Pharma) was injected locally at the site where the incision was made. The right masseter was partially removed and a large craniotomy (∼5 mm diameter) was performed above the auditory cortex using bone sutures of the skull as a landmark. For 2P calcium imaging, we did 3 to 5 injections at 200 μm intervals of 150 nL (25 nl/min) using pulled glass pipettes of rAAV1.syn.GCamP6s.WPRE virus (10^13^ virus particles per ml) diluted 30 times (Vector Core, Philadelphia, PA, USA). After this, the craniotomy was immediately sealed with a 5 mm circular coverslip using cyanolite glue and dental cement (Ortho-Jet, Lang) and a metal post for head fixation was implanted and fixed with dental cement on the top of the skull.

For labelling of thalamo-cortical fibers, rAAV1.syn.GCamP6s.WPRE virus (10^13^ virus particles per ml) undiluted virus was stereotaxically injected into auditory thalamus (MGN) (Anteroposterior = −3.0 mm, Lateral=2.1 mm, Dorsoventral=3.2). The fluorescence from thalamo-cortical fibers projecting into layer 1 of the auditory cortex could be recorded 4-5 weeks after injection.

### Two-photon imaging

For all isoflurane experiment, imaging was performed using a two-photon microscope (Femtonics, Budapest, Hungary) equipped with an 8 kHz resonant scanner combined with a pulsed laser (MaiTai-DS, SpectraPhysics, Santa Clara, CA, USA) tuned at 900-920 nm using a 10x objective (0.6 NA, XLPLN10XSVMP, Olympus, Japan) immersed in ultrasound transmission gel (Aspet, USA) previously centrifuged to eliminate air bubbles. Images were acquired at 31.5 Hz during blocks of 300 s. The imaging field of view was 1×1mm.

For ketamine-medetomidine and Zoletil® experiments, imaging was performed using an acousto-optic two-photon microscope (Kathala Systems, Paris, France) combined with a pulsed laser (Insight X3 Dual, SpectraPhysics, Santa Clara, CA, USA) tuned at 920 nm using a 16x objective (0.85 NA, Nikon, Japan) immersed in ultrasound transmission gel. The imaging field of view was 0.47×0.47mm, but four planes interspaced by 50 μm could be imaged simultaneously thanks to ultrafast defocusing with the acousto-optic deflectors. These four-plane stacks were imaged at 30.5 Hz during blocks of 150 s.

Mouse was habituated to stand still head fixed under the microscope during one week 30-60 min per day before the recordings. Recordings were performed inside a light-tight, sound proof box.

Imaging fields-of-view in the auditory cortex were 1×1mm. Imaging was performed at different depths ranging from 150 μm to 600 μm.

In 5 mice, after one awake imaging session, isoflurane anesthesia mixed in pure air was applied through a nose mask using the SomnoSuite anesthesia unit (Kent Scientific), without changing the field of view (example on field-of-view stability in **Fig 1a**,**b**). An infrared heat pad (Kent Scientific) was placed under the tube containing the mouse. 3% of isoflurane was applied for 1 min to induce narcosis. Anesthesia was then slowly decreased until we observed whisker movements and a level close to 1.3% was applied during recordings (actual range 1.2-1.4%). Typically, the limit of narcosis was observed between 0.9 and 1.1% isoflurane concentration based on the occurrence of spontaneous whisker movements which were monitored with an infrared camera (Smartek Vision, objective Fujinon/25mm). From this limit value, we increased concentration by 0.3%. Anesthesia was maintained at the same level during the 40 min of the imaging session.

Ketamine-medetomidine (KM) or Zoletil® (a 50-50% mix of tiletamine and zolazepam) were injected subcutaneously while leaving the animal under the microscope in head-fixation in order to maintain an identical field of view. We applied a dose of 50 mg/kg ketamine and 1 mg/kg medetomidine which after an induction of 10 min was sufficient to maintain the animal stably anesthetized for more than one hour. After the imaging session, atipamezole was injected intramuscularly to accelerate waking up. Zoletil® was applied at a dose of 70mg/kg (i.e. 35mg/kg tiletamine and 35mg/kg zolazepam). This dose was sufficient for maintenance of all 3 tested animals under anesthesia during 45min.

### Stimulation protocol and sounds

The following sequence was applied in awake state and then under anesthesia: two 300s recordings without auditory stimulation (pre-stimulation on-going activity), three 300 s recordings with sound stimulation then followed by two 300s recordings without stimulations (post stimulation on-going activity measurements). The same protocol was used under anesthesia (**Fig. 1a,b**). Each sound was 500 ms long and there was a one second interval between the end of one sound and the beginning of the next one. All sounds were delivered at 192 kHz with a NI-PCI-6221 card (National Instrument) driven by Elphy (G. Sadoc, UNIC, France) through an amplifier and high-frequency loudspeakers (SA1 and MF1-S, Tucker-Davis Technologies, Alachua, FL). Sounds were calibrated in intensity at the location of the mouse ear using a probe microphone (Bruel & Kjaer).

During each of 300s stimulations sessions, each sound was played 4 times in a random order chosen (in total 12 presentations for each sound). We used a set of predefined 50 sounds divided into 4 groups (**Extended Data Fig. 1**): 6 frequency modulated sounds (6 to 10, 10 to 16, 25 to 40 kHz, upward and downward modulations), 10 complex sounds, 6 pure tones (4, 6, 10, 16, 25, 40 kHz), 6 amplitude modulated sounds (sinusoidal modulation at 20, 7 and 3Hz, for three carrier frequencies of 25kHz and 4kHz). All sounds (except for sinusoidal) were played at two intensities of 60 and 80 dB SPL.

### Calcium signals processing, spike train estimation, neuropile traces

Data analysis was performed using Matlab scripts. Motion artifacts were first corrected frame by frame, using a rigid body registration algorithm. A single set of regions of interest (ROIs) corresponding to the neurons were defined by running Autocell (https://github.com/thomasdeneux/Autocell), a semi-automated hierarchical clustering algorithm based on pixel covariance over time ^17^, on the concatenated data from the awake and anesthesia sessions. Neuropil contamination was subtracted ^16^ by applying the following equation : F_corrected_ (t) = F_measured_ (t) – 0.7 F_neuropil_ (t), where F_neuropil_ (t) is estimated from the immediate surroundings (Gaussian smoothing kernel ^3^, excluding the ROIs, s = 170μm). The average neuropil signals shown in **Extended Data Fig. 7** are computed by averaging F_neuropil_ (t) across all putative neurons. We then applied to the neuropil corrected raw fluorescent signal the MLSpike deconvolution algorithm ^18^ (github.com/Mlspike) that finds the most likely spike train underlying the recorded fluorescence using a maximum-likelihood approach taking into account baseline fluorescence fluctuations. The parameters used by the algorithm were the typical time constant of calcium transient (1.7 s) and a coefficient (range used: 4 to 6) adjusting baseline drift compensation. Both of these parameters were estimated to best fit the descending slope of experimental calcium spikes as well as the fluctuation of the baseline fluorescence. Time constant estimation was in accordance with the published estimations for GCamp6s dynamics. After this process, every putative spike was described by its estimated onset time. We used the same spiking identification algorithm for the thalamo-cortical terminals with slightly different parameters (time constant 1.2 s and baseline drift compensation coefficient 6).

### Population events identification

After estimating spike trains with MLSpike (capable to estimate dense firing patterns (up to 20 Hz), where fluorescence rarely decays back to baseline), single cell activity was described as a binary vector, where the number of elements was equal to the number of time frames during the recording (frame duration was 31 ms). A value 1 was assigned at the time frames corresponding to the onset of each spike, and the vector was zero otherwise. The number of vectors was equal to the number of recorded neurons in the field of view, resulting in a matrix where columns corresponded to the time frames and rows to the neurons. Each column was summed to yield the number of neurons coactive at every time frame from which the instantaneous population firing rate could be deduced. As it is visible from sample raster plots (**Fig. 1** and **Extended Data Fig. 1**), there were periods of time where spikes were much more synchronized across the population, which could be interpreted as population events, departing from the fluctuations of an asynchronous population spiking process. To identify i) whether those peaks of activity were above asynchronous activity baseline and thus above by chance coincidence ii) when exactly the periods of synchronization started and ended, we applied the following algorithm:

1. The order of interspike intervals of each neuron was independently reshuffled 100 times, so that 100 surrogate matrices (number of neurons x number of time frames) were created. For each of these surrogate dataset, a population firing rate was calculated. A new matrix (100x number of time frames) where each row is a population firing rate for each of 100 reshuffling trials. From this matrix, we extract the 99 percentile of the surrogate distribution within each time frame. The average 99-percentile across time frames was then calculated and the final firing rate threshold was obtained by adding this average value to a local baseline estimate that aims to correct slow fluctuations of background firing rate. The baseline estimate was generated using asymmetric least squares smoothing algorithm with the parameters p=0.01 for asymmetry and λ=10^8^ for smoothness adapted from Eilers & Boelens, 2005^22^.
2. Experimental population firing rate trace was smoothed using Savitzky-Golay filtering (with order 3 and frame length 7) in Matlab to get rid of non-essential peaks (**Extended Data Fig. 1e**, gray trace). Local maxima for the smoothed curve that were above the previously defined threshold were retained (red dots). The adjacent local minima around each of the local maxima were identified (blue and green circles). Time frames identified this way were considered as the beginning (blued) and the end (green) of population events (**Extended Data Fig. 1e**). All the neurons that had at least one spike during this time interval were defined as a neuronal assembly.
3. Neuronal assemblies were then described as binary vectors of length equal to the total number of neurons in the recorded field of view. A value of 1 indicated the participation of the cell to the assembly with at least one spike over its duration and a value of zero indicated an absence of participation.

Neuronal assemblies could be detected in the same way during periods of stimulation and periods without stimulations (**Extended Data Fig. 1e**). However, in most of our analyses, neuronal responses evoked by sounds were quantified irrespective of the detection of a population event after sound onset. All the neurons that emitted a putative spike during sound presentation (within 500 ms interval from the beginning of the sound onset) were considered as belonging to an evoked neuronal assembly. All population responses to sounds were taken in account (12 per sound) except if zero spikes were detected across the entire population as occasionally happened during anesthesia.

### Clustering of assemblies

The Pearson’s correlation matrix for ongoing assemblies and sound-evoked responses was constructed by computing the Pearson correlation coefficient between binarized population vectors. For plotting the matrix, color intensity reflects the degree of correlation and the colormap was ranging from 0 to 1 unless otherwise stated. First, assemblies are arranged in a chronological order as shown in **Extended Data Fig. 1**. Then assemblies observed in the pre and post-stimulation blocks were separately reorganized using hierarchical clustering (agglomerative linkage clustering with furthest distance based on correlation metrics). Similar assemblies (i.e. that shared a substantial number of neurons), were clustered together (**Fig 1, Extended Data Fig. 1**). For evoked responses, the organization was simply based on sound identity (except in **Extended Data Fig. 2** where clustering was also applied to actually detected assemblies). In this case, the size of each group was constant as every sound was presented 12 times during an imaging session.

To quantify the reproducibility of same sound responses or of ongoing assemblies within clusters, correlation was calculated across all sound repetitions or across all assemblies of a cluster. Similarity across clusters and sound responses was calculated as the average correlation between all pairwise elements of the two compared groups of assemblies/responses. To measure maximal similarity between spontaneous and evoked activity, for each group of evoked response, the spontaneous cluster with maximum cross-correlation was identified.

For a given sound *i*, we estimate the similarity of population responses to on-going populations events as:

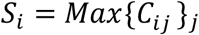

where *C*_*ij*_ = < *r*_*ij*_(*k, l*) >_*k,l*_ and *r*_*i j*_(*k, l*) is the Pearson correlation coefficient between population vectors observed for presentation *k* (range 1 to 12) of sound *i* (range 1 to 50), and for assembly*l* from the on-going assembly cluster number *j. Max*{*C*_*ij*_}_*j*_is the maximum *C*_*ij*_ across all spontaneous clusters (i.e. mean correlation with the on-going event cluster *j*_*i*_ that is the most similar to sound *i*).

The similarity was compared to the mean internal reproducibility of responses to sound *i* and of assemblies within clusters *j*_*i*_:

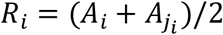

where 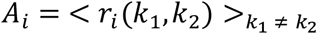 is the average correlation across all distinct pairs of responses to sound *i* and 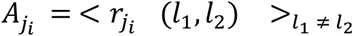 is the average correlation across all distinct pairs of on-going assemblies in cluster *j*_*i*_ which has maximal similarity with responses to sound *i. r*_*i*_ (*k*_1_, *k*_2_) (respectively 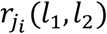 is the correlation between distinct pairs of population vectors indexed by *k*_1_and *k*_2_ (range 1 to 12) observed for sound *i* (respectively indexed *l*_1_and *l*_2_ for on-going activity clusters *j*_*i*_). If sound-evoked assemblies are similar to a cluster of events occurring spontaneously, we expect *S*_*i*_ ≈ *R*_*i*_, otherwise *S*_*i*_ < *R*_*i*_.

### Dimensionality reduction

The neuronal state space is defined as the space of activity (probability to respond to a sound or participate in an on-going assembly) of all simultaneously recorded neurons. A given event is represented by a vector that has the dimension of the neuronal population. To represent population vectors in a three dimensional space that capture a maximum of the variance without imposing non-linear distortions of the space, we used principal component analysis (PCA). PCA was performed on a dataset pulling all on-going assemblies and sound responses during both anesthesia and wakefulness. Projection of population vectors on the first three principal components were used for displaying them in a 3-dimensional space as in **Fig. 3**.

### Neuronal response probability distributions, sparseness and probability models

Lifetime sparseness was measured through the kurtosis of the response distribution ^27^, a measure that is more general than other sparseness measures as it applies also to negative responses.

Every neuron was characterized by a probability to respond to any sound (abscissa) and to participate in any spontaneous population event (ordinate) (**Extended Data Fig. 9a**). The neurons plotted in this space were divided into 3 groups separated based on the mean absolute deviation (MAD) of the difference between evoked and spontaneous probabilities. All the neurons for which the probability difference was smaller than 1 MAD and larger than −1 MAD were classified as neurons with equal probability to respond to sounds or to participate in a spontaneous event (**Fig. 6a**, green). Those with a negative probability difference below 1 MAD (**Fig. 6a**, red) were classified as having higher probability to respond to sounds, and those with a positive difference above 1 MAD (**Fig. 6a**, blue) were classified as having higher probability to participate in spontaneous events. Mean absolute deviation was chosen instead of standard deviation because the probability difference was not normally distributed.

To better capture how responsiveness to on-going and evoked events were distributed across the population, we defined 3 different probability models. The first model simply assumes that the probability to be active in an ongoing event is independent of the probability to be active in an evoked response. The expected distribution of probabilities for this hypothesis was generated by randomly shuffling observed probabilities across cells, generating a distribution that is much broader than the one observed for our data. The second model assumes that the probability to be active in an on-going event and in a sound response is identical. This model was simulated by defining first the distribution of probabilities to participate in an ongoing event for 6310 surrogate neurons. The average probability is set to 0.11 ± 0.08, in accordance with experimental data. Identical values were attributed to the probabilities to respond to any sound.

To more closely simulate the wakefulness condition, the last model supposes that for every neuron *i*, the probability of response to a sound is *p*_*resp*_*(i) = p*_*common*_*(i)+p*_*sound*_*_*_*spec*_*(i)*, and the probability of participation to an on-going event is *p*_*ongoing*_*(i) = p*_*common*_*(i)+ p*_*spont*_*_*_*spec*_*(i)*. The value of *p*_*common*_*(i)* is drawn from a random distribution *exp(-(x/k)*^*2*^*)/S* for *x* between 0 and 1. *S* is the integral of *exp(-(x/k)*^*2*^) between 0 and 1. *p*_*sound*_*_*_*spec*_*(i)* and *p*_*spont*_*_*_*spec*_*(i)* are drawn from to independent Gaussian distributions centered on 0 and of variance *v*^*2*^_*sound*_ and *v*^*2*^_*spont*_. The parameters were chosen to fit experimental distribution of probabilities difference, *k* = 0.1, *v*_*sound*_ *= 0*.*85* and *v*_*spont*_*= 0*.*65*.

### Template matching classifier

To quantify the sound specificity of the patterns of neuronal assemblies, we used a cross-validated template matching algorithm where correlation was the metric between population vectors. Every sound was presented 12 times. We used a leave-one-out cross-validation procedure with a training set of 11 sound presentations and a test set of 1 sound presentation. This was repeated 12 times, changing the test sound presentation each time. At every iteration of classification, the response to the test sound presentation was compared to the 50×12 - 1 other single trial sound responses using the correlation distance as a metric. The test response was then attributed to the sound for which it had the smallest average distance with its single-trial responses (excluding the test response in the calculation).

### Clustering of single neurons

Clustering was also used to organize neurons according to the similarity of their responses to the sounds. Due to the large variability observed in many neurons, this analysis is not exhaustive but rather aims at identifying principal classes of responses within our dataset. Clustering was performed across the 5 cortical imaging sessions thus including (3641 neurons) and the 7 thalamic axon imaging sessions (13314 terminals), in both awake and anesthetized states. Each neuron was characterized by a vector of 50 elements (corresponding to the number of presented sounds), where each element contained the information about the number of responses during the trial (from 0 to 12). A neuron was considered to respond to a sound if it generated at least 1 spike during 500 ms of sound duration. The Pearson’s correlation matrix for all neuron/terminal response vectors was constructed by computing the Pearson correlation coefficient between them. Then neurons/terminals were reorganized using hierarchical clustering (agglomerative linkage clustering with furthest distance based on correlation metrics). The groups of neurons sharing similar sound response profiles were assessed. This method yielded in a number of strongly correlated clusters of neurons that were tuned to multiple sounds, few clusters specific to a single sound and several small clusters, which after visual inspection appeared to contain noisy responses (hence very dissimilar to other clusters). Several of the clusters of thalamo-cortical terminals were single-sound specific. In case of thalamo-cortical terminals, due to the response sparseness across the large population putative axonal terminals, only the ones that responded to any sound at least twice were taken into consideration before clustering (representing 25% in awake state and 5% under anesthesia). To identify the sounds to which every cluster was significantly tuned at different conditions (awake and anesthesia) we calculate, the sound responses of every neuron within a given cluster were reshuffled over all the stimulation instances (597). The number of non-zero reshuffled responses for 12 randomly chosen instances (out of 597, corresponding to the number of presentations of the same sound during the trial) was averaged over all neurons of the cluster. We used one-way Anova (Matlab, The Mathworks), separately applied to every cluster and every condition (awake, anesthesia), to identify the sounds that had significantly higher responses than expected from the shuffled dataset. The Anova function also returned the particular sounds for which the response was significant.

### Statistical analysis

All statistical analyses were performed using built-in MATLAB functions unless otherwise noted. Exact P values for every statistical test performed are mentioned in the figure captions. For all box-and-whiskers plots, the red mark indicates the median, and the bottom and top edges of the box indicate the 25th and 75th percentiles, respectively. The whiskers extend to the most extreme data points not considered outliers, and the outliers are plotted individually using the ‘+’ marker symbol. The following tests (precised in every figure caption) were used: One-way analysis of variance (ANOVA) followed by Tukey’s multiple comparison test: anova1 and multcompare, Wilcoxon Rank Sum Test (two-sided, the equivalent of the Mann-Whitney U test), Paired Wilcoxon Signed Rank Test (two-sided).

## Supporting information

Exetended data

## Acknowledgements

We thank Thomas Deneux, Guillaume Hucher and Aurélie Daret for their technical support, Melody Torao-Angosto for help with pilot experiments and numerous colleagues for comments on analyses and figures.

## Funding

European Community, Future and Emerging Technologies program (Human Brain Project, H2020-945539) (AD)

Paris-Saclay University “Initiatives de Recherches Stratégiques” -NeuroSaclay and Icode (BB, AD)

Agence Nationale pour la Recherche 12-PDOC-0006, (BB), PARADOX (AD) Région Ile de France - DIM Cerveau Pensée - MULTISENSE (BB)

Fondation pour l’Audition, FPA IDA02 and APA 2016-03 (BB). European Research Council, ERC CoG 770841 DEEPEN, (BB)

BB acknowledges the support of the Fondation pour l’Audition to the Institut de l’Audition.

## Author contributions

A.F, A.D and B.B. conceived experiments, designed the study and interpreted data. A.F. and J.S. collected data and A.F. and B.B. performed the analyses. A.F, A.D. and B.B. prepared figures and wrote the manuscript. A.D. and B.B. managed the project.

## Data and materials availability

All data used in the analysis are available on Zenodo (10.5281/zenodo.5270513), code and materials are available upon request.

## Extended Data

Extended Data Figs. 1 to 9

