## Supplementary material for "Awake perception is associated with dedicated neuronal assemblies in cerebral cortex": Exetended data

for

**This PDF file includes:**
Extended Data Figs. 1 to 9

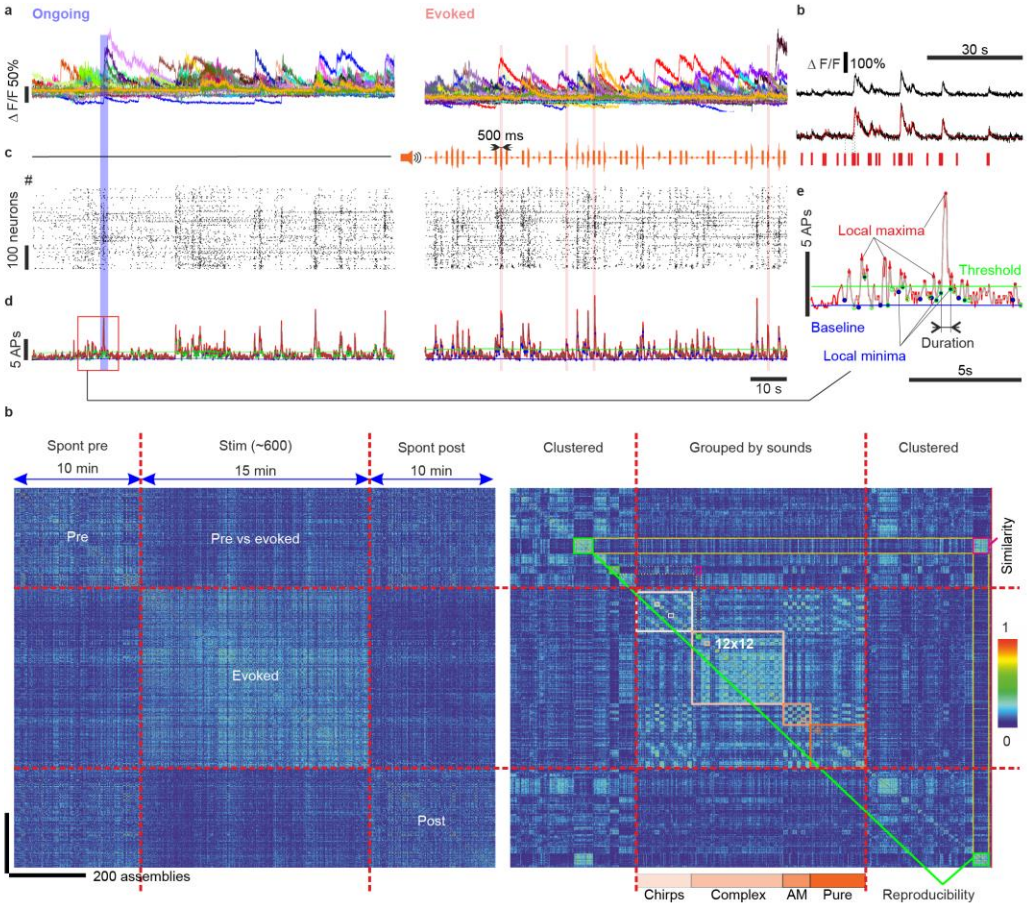


**Extended Data Fig. 1: Identification and clustering of spontaneous and evoked assemblies.**
**(a)** Examples of raw calcium fluorescence traces without stimulation (left) and during sound stimulation (right). **(b)** Spikes time estimates (red) were extracted from calcium fluorescence traces

(blue) using the MLSpike algorithm (green trace is the raw fluorescent trace adjusted to the baseline fluorescence fluctuations). **(c)** Population raster plots during no stimulation (left) and stimulation (right) periods. Every dot represents a spike estimated as in B. **(d)** Population firing rate (30ms bins) during no stimulation (left) and stimulation (right) periods. Vertical transparent bars highlight spontaneous (blue) and evoked (pink) population events detected as described in E (inset). **(e)** Identification of spontaneous population events from population firing rate (red trace). Stochastic threshold (green), based on population spiking activity and added to the baseline (blue), to identify the local maxima above the threshold (red dots) in the smoothed population firing rate trace (grey). Local minima (blue and green dots) adjacent to the above threshold local maxima define the beginning and the end of the population event, respectively. All neurons spiking at least once within this interval are included into the population event, defining spontaneous or evoked neural assemblies. **(f)** Neuronal assemblies, where the length of the binary vector is the total number of neurons in the FOV, and the 1 elements correspond to the neurons active during the respective population event (in red). **(g)** For an example recording session, Pearson's correlation matrix between spontaneous assemblies (pre- and post-stimulation) and the assemblies evoked by a single trial sound response (50 sounds presented 12 times each) in the order of their appearance during 35-minute recording in awake mouse. Diagonal quadrants show the auto-correlation between the same types of assemblies, lateral quadrants – cross-correlation between different types of assemblies **(h) Upper panel**: the types of sounds used for stimulation. **Lower panel**: same matrix as in G where spontaneous assemblies are sorted by hierarchical clustering and evoked responses (whether or not a population event was detected) are sorted sound-by-sound (12 trials/sound). The mean correlation among the assemblies in a cluster or among the assemblies corresponding to single trial sound responses define the reproducibility of a spontaneous or evoked pattern (orange inset). The mean cross-correlation between two groups of patterns defines their similarity (purple inset). Color scale for pattern correlation values (red for 1, dark blue for 0). **(i)** Spectrograms of complex sounds (each of them have a duration of 500 ms and are presented at 60 and 80dB intensities).

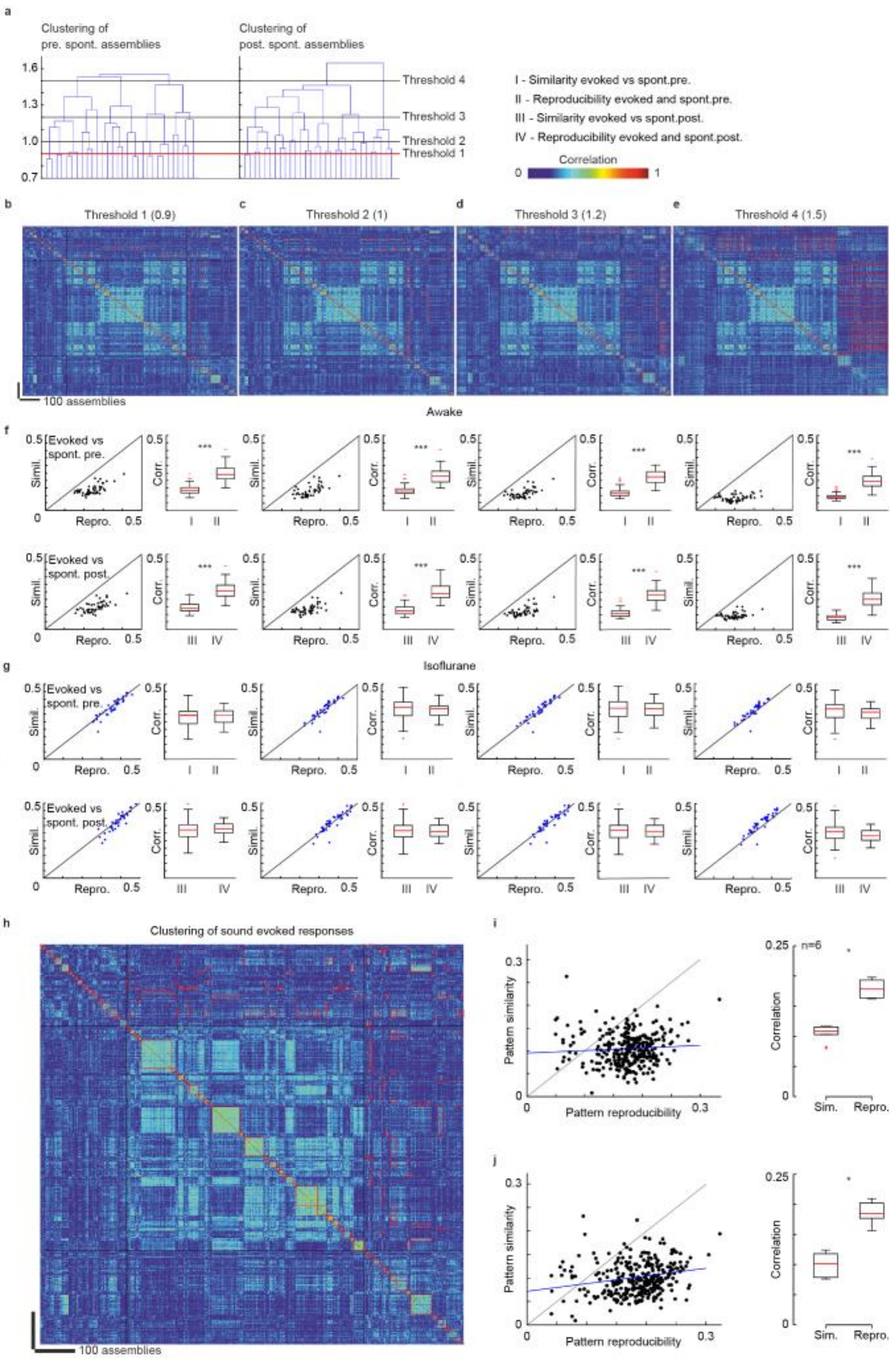


**Extended Data Fig. 2: Dissimilarity of evoked and on-going assemblies is robust to clustering methods and parameters.**

**(a)** Dendrogram plot of the hierarchical binary cluster trees for spontaneous assemblies before and after stimulation in a sample recording session. The horizontal lines indicate the 4 cophenetic distance thresholds tested in the following panels (0.9, 1, 1.2, 1.5). **(b-e) Upper panel**: Correlation matrix for a sample recording in the awake state where spontaneous assemblies are sorted by hierarchical clustering with thresholds 1, 2, 3 and 4 (in **b**, **c**, **d** and **e** respectively) as defined in A. Red squares in the lateral quadrants indicate the maximum similarity pairs. Red squares in the diagonal quadrants indicate clusters and groups. (**f)** Scatter plot and statistics of similarity vs reproducibility (evoked/spont. pre., top; evoked/spont.post., bottom) for each threshold value for the example shown in **b-e**. Reproducibility of spontaneous and evoked assemblies is significantly higher than the similarity of evoked vs spontaneous assemblies independently of the threshold ($p<0.0001$, across 50 pairs of clusters, Paired Wilcoxon Signed Rank Test). **(g)** Same as **f** but for the anesthetized state under isoflurane in the same sample population ($p>0.05$, across 50 pairs of clusters, Paired Wilcoxon Signed Rank Test, thresholds: 0.6, 0.8, 1 and 1.2). **(h)** Sample matrix where evoked assemblies were clustered in the same way as spontaneous ones (instead of grouping by sound). **(i)** Relation between reproducibility (abscissa) and similarity (ordinate) of sound-evoked and spontaneous pre-stimulation assemblies for all sounds and sessions. Statistics across sessions are given in the right-hand-side histograms ($p=0.03$, $n=6$ mice). **(j)** Same as in **i** for evoked *versus* spontaneous assemblies detected post-stimulation ($p=0.03$, $n=6$ mice). Paired Wilcoxon Signed Rank Test for **i** and **j**.

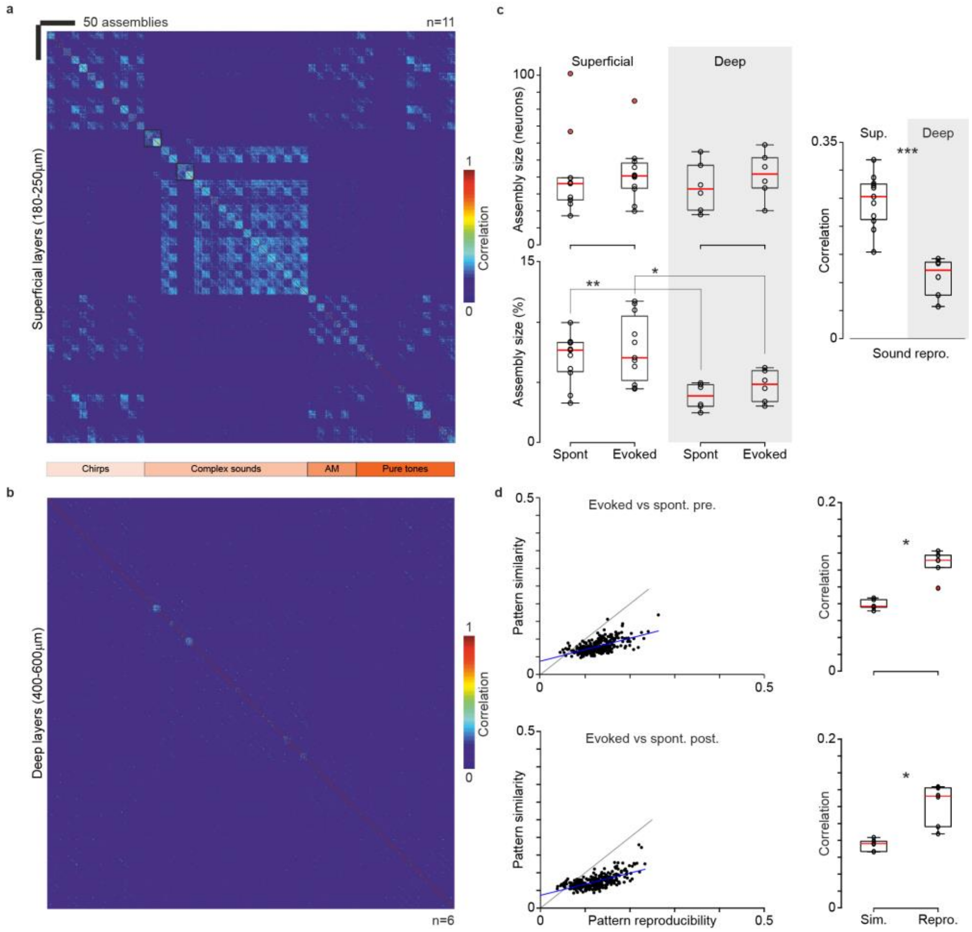


**Extended Data Fig. 3: Dissimilarity of evoked and on-going assemblies is seen also in deep cortical layers**

**(a)** Pearsons's correlation matrix averaged across 11 mice for the assemblies evoked by sounds in the upper layers of auditory cortex (180-250 um below cortical surface). **(b)** The same as in A, in deep layers (450-600 um below the cortical surface) averaged across 6 mice. **(c)** Top left: average size (number of neurons) of neuronal assemblies in superficial (11 mice) and deep layers (6 mice). Bottom left: average size of neuronal assemblies in % of all neurons in the FOV in superficial and deep layers (p=0.005, p=0.02, Wilcoxon rank sum test). Right: sound evoked responses reproducibility in superficial layers is higher than in the infragranular layers (p=0.0002, Wilcoxon rank sum test). **(d)** Dissimilarity of sound-evoked and spontaneous patterns for all sounds and sessions in deep layers. Statistics across sessions are given on the right-hand-side histograms (p=0.03, p=0.03, Paired Wilcoxon Signed Rank Test, n=6 mice).

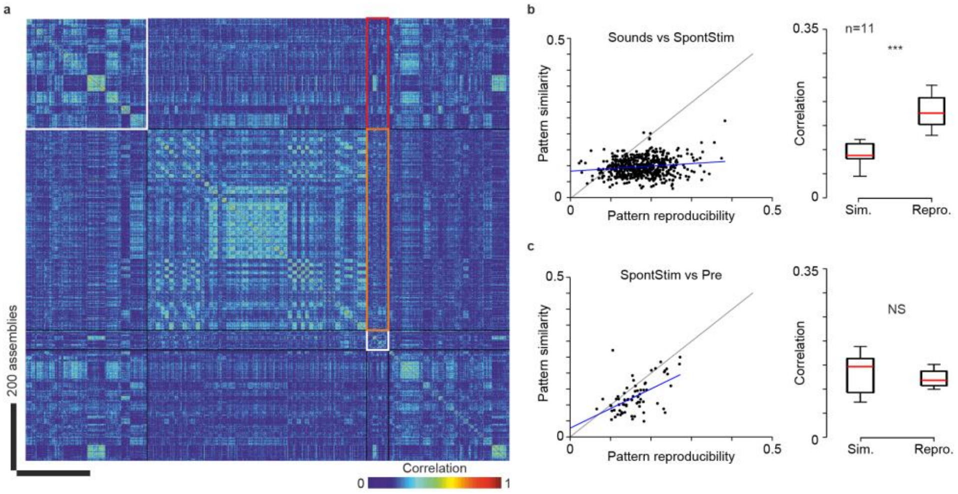


**Extended Data Fig. 4: On-going assemblies during stimulation are also dissimilar to evoked responses**

**(a)** For an example recording session, Pearson's correlation matrix between spontaneous assemblies sorted by hierarchical clustering and single trial sound response patterns. Spontaneous assemblies detected during stimulation sessions in between evoked responses are clustered in the small white rectangle. Their similarity with evoked and spontaneous pre-stimulation assemblies (big white rectangle) are within orange and red rectangles, respectively. **(b)** Dissimilarity of sound-evoked and spontaneous patterns (SpontStim) detected during stimulation (reproducibility on abscissa and similarity on ordinate). Statistics across sessions are given on the right-hand-side histograms ($p=0.001$). **(c)** Similarity of on-going assemblies observed during the pre-stimulation and stimulation epochs. ($p=0.41$, Paired Wilcoxon Signed Rank Test, $n=11$ mice in B and C).

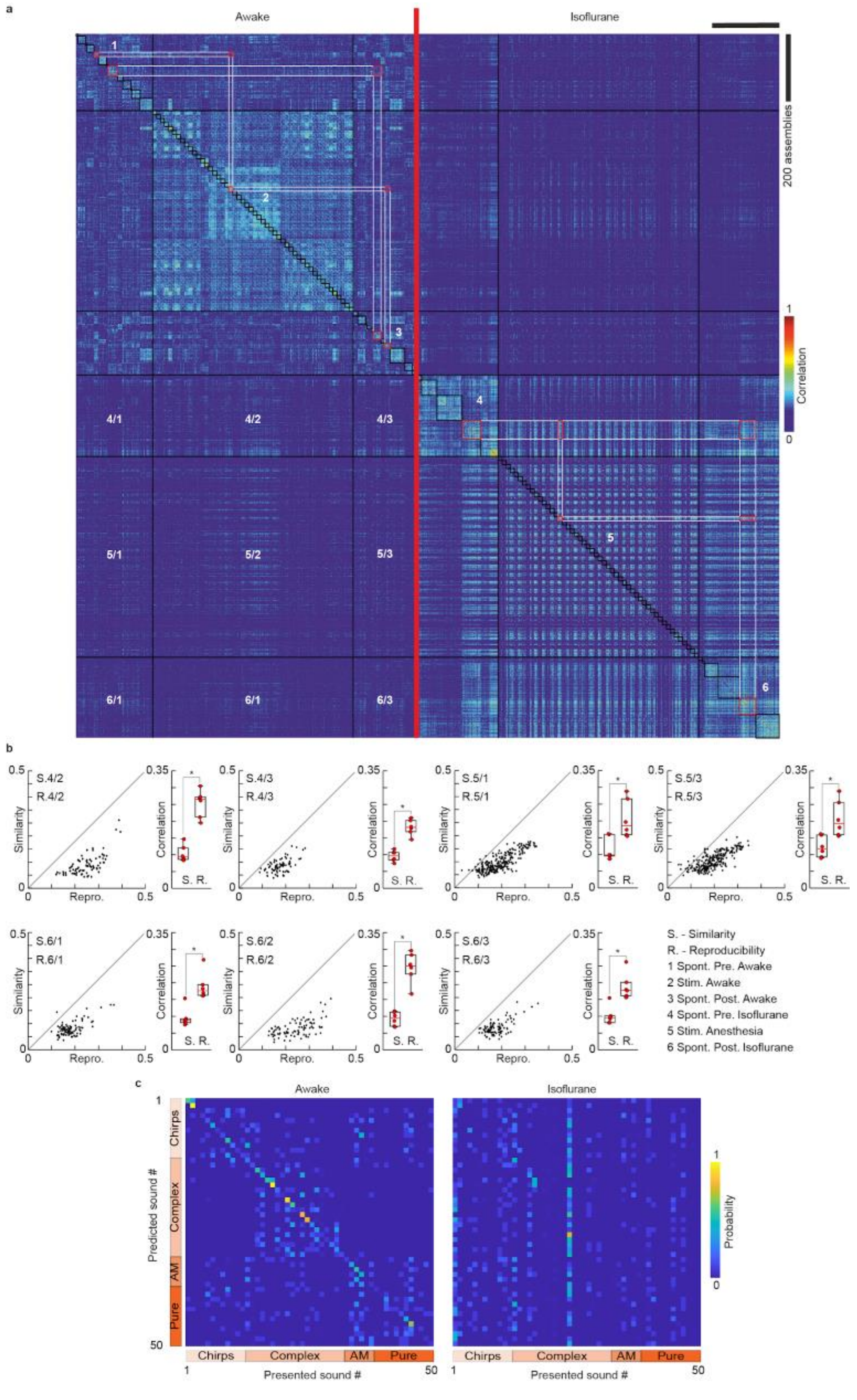
a
Awake
Isoflurane
200 assemblies
Correlation
1
0
1
2
3
4
5
6
4/1
4/2
4/3
5/1
5/2
5/3
6/1
6/1
6/3
b
S.4/2
R.4/2
S.4/3
R.4/3
S.5/1
R.5/1
S.5/3
R.5/3
S.6/1
R.6/1
S.6/2
R.6/2
S.6/3
R.6/3
Similarity
Correlation
Repro.
S. R.
S. - Similarity
R. - Reproducibility
1 Spont. Pre. Awake
2 Stim. Awake
3 Spont. Post. Awake
4 Spont. Pre. Isoflurane
5 Stim. Anesthesia
6 Spont. Post. Isoflurane
c
Awake
Isoflurane
Predicted sound #
Presented sound #
Chirps
Complex
AM
Pure
Probability
1
50
0

**Extended Data Fig. 5: Transition from awake to anesthesia changes both sound-evoked and spontaneous patterns.**

**(a)** Correlation matrices for on-going and evoked assemblies across wakefulness and anesthesia. The numbering of the quadrant describes the different recording conditions. (1) On-going assemblies in the awake state, sorted by hierarchical clustering (black rectangles). (2) Single trial sound responses in the awake state, grouped sound-by-sound (black rectangles). (3) Post-stimulation on_going assemblies in the awake state. (4) Ongoing assemblies under anesthesia, before stimulation. (5) single trial sound responses under anesthesia (whether or not a population event was detected) (6) Ongoing assemblies under anesthesia, after stimulation. Red rectangles in the top right quadrant outline the similarity between the assemblies detected in the awake and anesthetized states. Numbering in the bottom left quadrant indicates pairwise comparisons (e.g. 4/1 Spont. Pre. Anesthetized vs Spont. Pre. Awake). **(b)** Plot of awake versus anesthetized assembly similarity against assembly reproducibility. Statistics across sessions are given on the right-hand-side histograms (p=0.03, Paired Wilcoxon Signed Rank Test, n=6 mice). (**c**) Confusion matrix (i.e. probability ) of the classifier used in **Fig. 4e** averaged across the 5 mice in which the same populations were imaged in the awake and isoflurane anesthetized state.

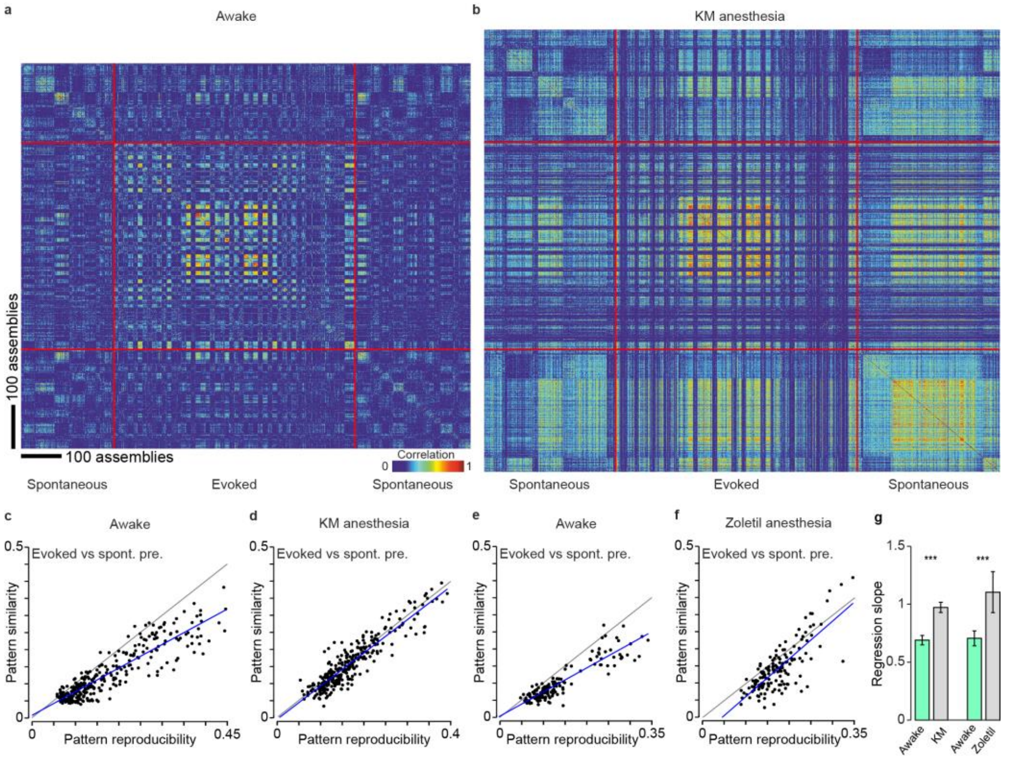


**Extended Data Fig. 6: Ongoing assemblies and sound-evoked responses overlap under ketamine-medetomidine and zolazepam-tiletamine anesthesia. (a)** For an example recording session, Pearson's correlation matrix between spontaneous assemblies sorted by hierarchical clustering and single trial sound response patterns (whether or not a population event was detected), sorted sound by sound (12 trials/sound). Clustering is done independently in pre- and post-stimulation periods. Lower correlation inside black and orange frames (similarity) compared to correlations along the diagonal (reproducibility) indicate that spontaneous and evoked patterns are different. **(b)** Correlation matrix under ketamine (50 mg/kg) medetomidine (1 mg/kg) anesthesia (KM) for the same neuronal population as in **a**. Similar correlation in black and orange frames (similarity) and in the squares along the diagonal (reproducibility) indicate that spontaneous and evoked assemblies are highly similar. **(c)** Relation between reproducibility (abscissa) and similarity (ordinate) of sound-evoked and spontaneous patterns for the 50 sounds and 6 sessions recorded in 3 mice. The blue line is a linear regression and the black line is the unity line. (**d**) Same as (c) but under KM anesthesia. (**e-f**) Same as a (c-d) but for 3 experiments in 3 different mice for Zoletil® (70mg/kg) anesthesia. (**g**) Slope of the regression lines in (c-f). Error bars indicate the 5% confidence interval obtained by bootstrapping across data points. Bootstrapping (1000 random resampling with replacement) was also used to assess the significance of the difference between slopes obtained in anesthesia and in wakefulness. For both anesthetics, the bootstrap p-value was <0.001. For KM anesthesia, the difference between awake and anesthetized was also significant when testing across the 6 sessions (Wilcoxon rank sum test, p=0.03). Note that the higher slopes observed here in the awake state compared to the results shown in Fig. 2 are likely due to the smaller field of views used in KM and Zoletil® experiments (see Methods), which reduced the variety of assembly configurations and therefore the distance between assemblies.

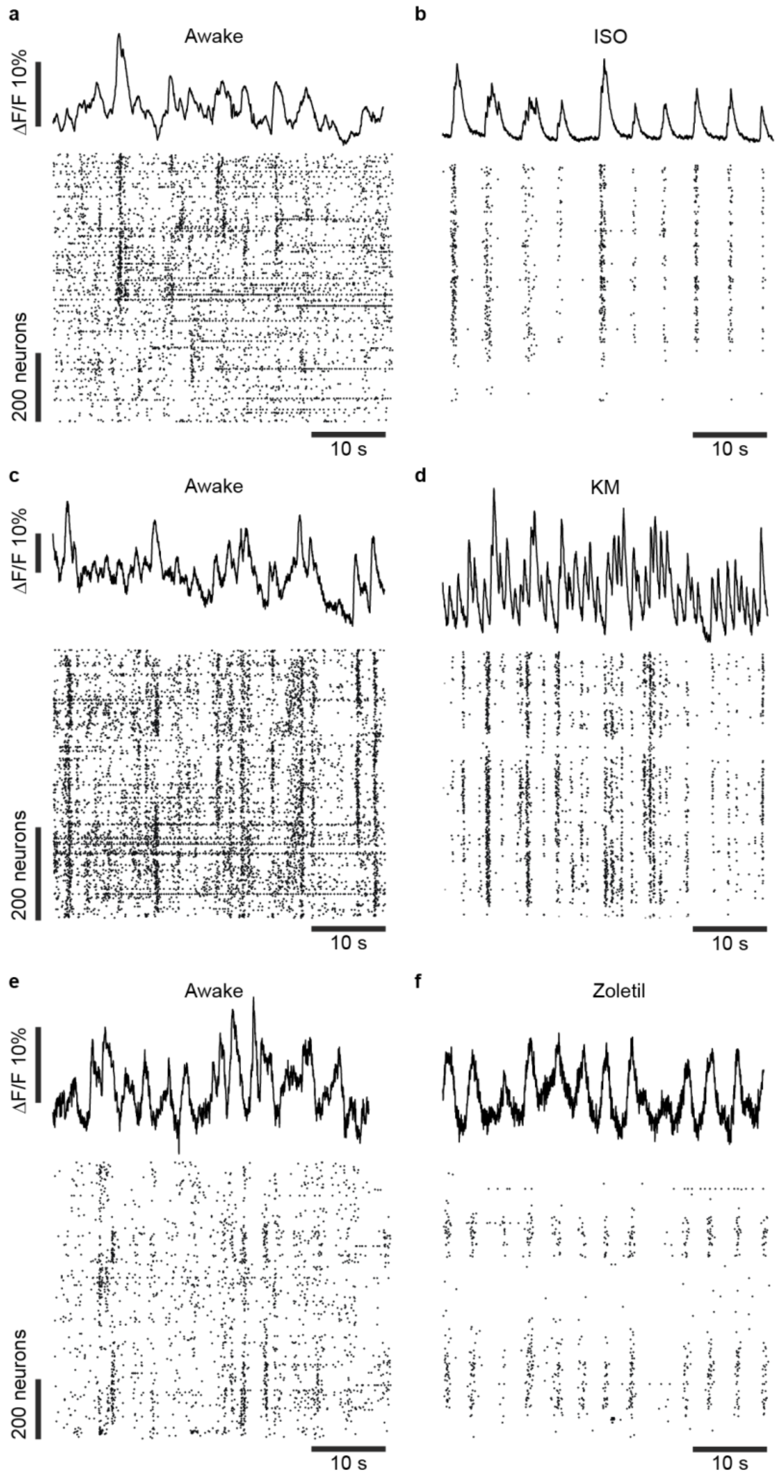


**Extended Data Fig. 7: Sample neuropil traces during wakefulness and anesthesia. (a) top panel** Sample trace of the mean neuropil signal for a representative imaging session during the awake state. Neuropil was measured in a disk of 20 pixel radius around each region of interest identified as neurons, excluding other neurons. **bottom panel** Raster plot of putative action potentials (MLSpike) in the recorded population align with the neuropil trace above. **(b)** Same as

(a) under isoflurane anesthesia. (**c-d**) Same as (a-b) but for ketamine medetomidine anesthesia. (**e-f**) Same as (a-b) but for Zoletil anesthesia.

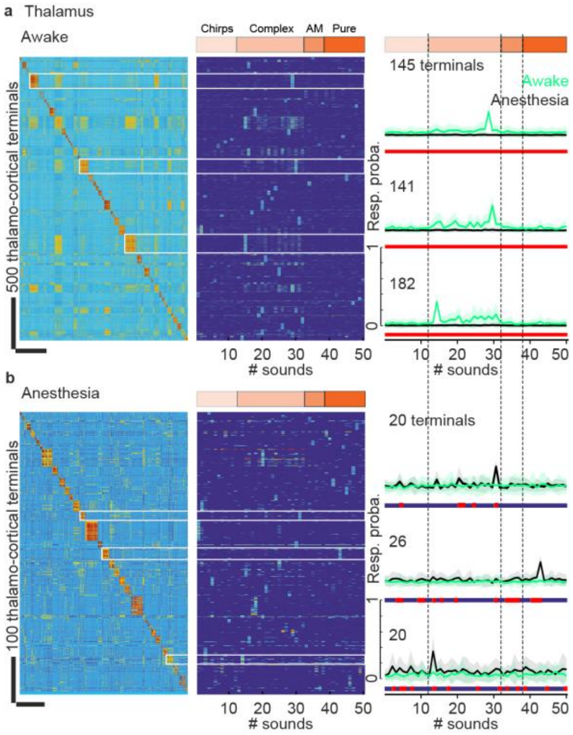


**Extended Data Fig. 8: Transformation of thalamic sound tuning profiles between wakefulness and anesthesia. (a) Left panel:** a fragment of response profile correlation matrix containing 2800 (out of 13314) clustered responsive thalamo-cortical terminals (7 mice) recorded in the layer 1 of the auditory cortex in the awake state. **Middle panel**: Response probability for thalamo-cortical terminals organized with the same clustering as for the matrix of the left panel (colours in the upper band indicate the type of sound). **Right panel**: Mean response profiles of thalamo-cortical terminals in the awake state (red traces) and under anesthesia (green trace) for the 3 sample clusters labelled in the left panel. **(b) Left panel:** a fragment of response profile correlation matrix containing 620 (out of 13314) responsive thalamo-cortical terminals (the same as in **a**) recorded under anesthesia and reclustered. **Middle panel**: Response probability for the thalamo-cortical terminals ordered according to the clusters in the left panel. **Right panel**: Mean response profiles of thalamo-cortical terminals under anesthesia (green trace) compared to their profiles in the awake state (red traces) for the 3 sample clusters labeled in the left panel. As for the cortical neurons, tuning profiles of thalamo-cortical fibers are significantly modified (Wilcoxon rank sum test, $p<0.05$) when passing from awake to anesthetized state.

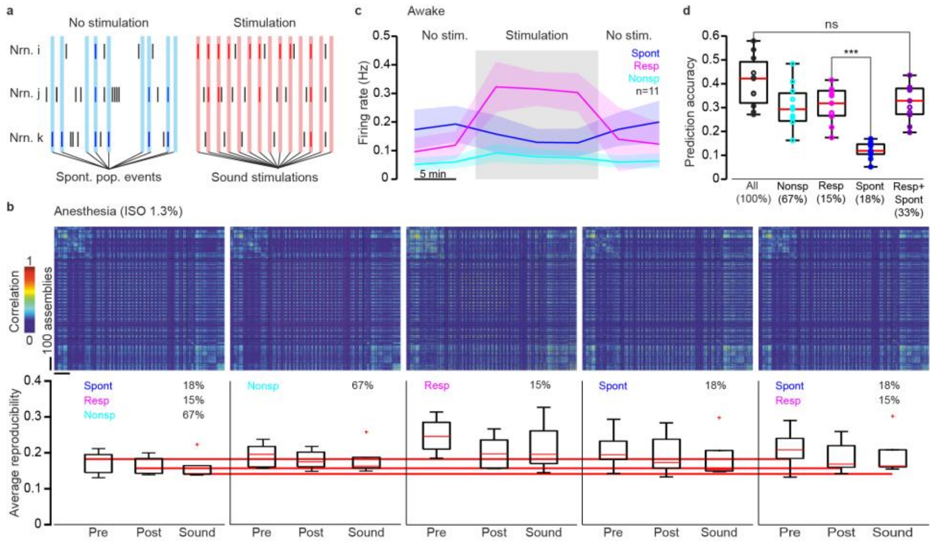


**Extended Data Fig. 9: Neuronal assemblies lose their functional specificity under anesthesia.**

**(a)** Schematics depicting the rationale for the selection of the three functional populations defined in **Fig. 6**. For each neuron the probability to participate in a spontaneous event (blue bars) and in a sound evoked event (red bars) is calculated. In the evoked *versus* spontaneous probability space (right panel), blue dots represent neurons with higher probability to be activated spontaneously, than to be activated by sound. Green dots represent neurons with a similar probability to be engaged in both kinds of activity; red dots are neurons more probably activated by a sound. **(b) Top left:** For a sample session under anesthesia, Pearson's correlation matrix computed with all available neurons. **Bottom left**: Average sound and spontaneous event cluster reproducibility under anesthesia. The neurons are divided into 3 groups based on their activity in awake state before anesthesia application. **From left to the right**: Same as the first matrix but only for Nonspecific neurons; only for predominantly sound responsive neurons (RESP); only predominantly spontaneously active neurons (SPONT); only for RESP+SPONT. No difference with each subpanel is significant (Wilcoxon rank sum test, 6 mice, $p>0.05$). **(c)** Time course of the mean firing rate profiles for each of the 3 groups of neurons in all awake mice (n=11). Note that the graph presented in **Fig. 7b** corresponds to only 5 mice that were recorded both in awake and anesthetized conditions. **(d) Left panel**: From top to bottom: Sound prediction accuracy taking into account only Nonspecific neurons, only sound responsive neurons (RESP), only predominantly spontaneous neurons (SPONT) and responsive and spontaneous together (RESP+SPONT) in awake state, n=11 mice. **(e)** Average prediction accuracy for all sounds for the 4 sub-populations (n=11 mice, $p=0.11$, $p=0.00001$, One-way ANOVA test).
